# Multi-level orchestration of γδ T cell development and central nervous system inflammation by retinoic acid

**DOI:** 10.1101/2020.08.12.247510

**Authors:** Marcelo Gregorio Filho Fares da Silva, Rasmus Agerholm, Nital Sumaria, John Rizk, Julia Fiske, Elisa Catafál-Tardos, Maria Virginia Baglioni, Davide Secci, Paula Peñalver Sebastián, Daniel Pennington, Vasileios Bekiaris

## Abstract

The vitamin A metabolite retinoic acid (RA) is critical for the maturation and function of the immune system, however, our knowledge regarding its role in gamma delta (γδ) T cells is limited. By specifically inactivating the RA receptor alpha (RARα) in type 3 lymphocytes and using a combination of fetal thymic organ cultures and single-cell RNA-sequencing we found that RA strengthened TCR signaling to limit the embryonic development of interleukin(IL)-17-producing γδ T cells (γδT17), while at the same time it promoted their survival after birth. In adult mice, RA imposed a multi-level transcriptional program allowing γδT17 cells to become activated, survive and sense interferon. Consequently, cells with inactive RARα responded poorly to immunization and produced low level of cytokines leading to near disease resistance in the experimental autoimmune encephalomyelitis model. Finally, RA was required for optimal expression of the α4β1 integrin and γδT17 cell infiltration to the brain. Collectively, these studies suggest that by acting at various stages during the lifetime of γδ T cells, RA supports their normal development, their survival and their ability to respond to inflammatory stimuli.

## Introduction

Gamma delta (γδ) T cells are innate-like lymphocytes, evolutionary conserved from lampreys to humans (*1, 2*). They are critical for effector immunity against many pathogens and cancers, but also have key tissue homeostatic functions, while in chronic inflammation they can become highly pathogenic (*3*). Similar to other lymphocytes, γδ T cells can be defined in subsets based on the type of immune responses they participate in and elicit, such as type 1 interferon(IFN)-γ-producing (γδT1) or type 3 interleukin(IL)-17-producing (γδT17) (*4*). Mouse γδT17 cells start to develop around embryonic day 16 (e16) through a combination of signals from the T cell receptor (TCR) and the transcription factors cMAF and RORγt. Thus, γδ T cell committed progenitors that have avoided Skint-1 selection (*5*), require weak TCR signals (*6, 7*) to differentiate into cMAF- expressing cells, a step important for their commitment into fully functional RORγt^+^ type 3 effector cells (*8*). Right before and right after birth γδT17 cells are exported from the thymus to secondary lymphoid organs (SLOs) and most epithelial and barrier surfaces (*9*). During the first week of life STAT5 is required to sustain γδT17 survival and proliferation within tissues (*10*), while continuous suppression of the non-canonical NF-κB pathway by the E3 ligases cIAP1 and cIAP2 maintains expression of cMAF and RORγt and allows further expansion after weaning (*11*). Thus, sequential embryonic and neonatal checkpoints ensure the lineage stability of γδT17 cells.

In addition to their seemingly pre-programmed phenotype and function, γδT17 cells acquire tissue-specific properties, such as type 1 effector functions in the gut (*10, 12*), or thermoregulation in adipose tissue (*13*). In muscle, γδT17 cells promote tissue regeneration following injury (*14*) and in the brain they appear to be important for maintaining short-term memory (*15*) and to regulate anxiety-like behavior (*16*). Collectively, these data suggest that γδT17 cells can sense and adjust to their microenvironment, likely by responding to local metabolites. In this regard, we reasoned that dietary associated metabolic products, such as vitamin derivatives, could be critical for the development and functional maturation thereafter of γδT17 cells.

Vitamin A is a natural retinoid that can only be obtained through the diet and is necessary for normal human development (*17*). Pre-formed dietary retinoids, mainly retinol and retinyl esters, or proretinoids like β-carotene are absorbed in the gut and after a series of hydrolysis and esterification reactions are packed into chylomicrons, and through the lymphatic system are primarily transported to the liver where they are stored (*18*). Retinol packed in the liver is transported to peripheral tissues via retinol binding proteins and upon cellular uptake it can get oxidized by aldehyde dehydrogenases into its main active metabolite retinoic acid (RA) (*18, 19*). Cellular RA binding proteins relocate RA from the cytosol to the nucleus where it will interact with RA receptor (RAR)-retinoid X receptor (RXR) dimers (*18, 20*). The RAR-RXR complex may be composed of any of three RARs, RARα, RARβ, RARγ, which are mostly ubiquitously expressed, but can show tissue specificity (*21, 22*). The RAR-RXR dimer functions as a transcriptional regulator, which upon binding to RA releases its associated co-repressors, recruits co-activators and initiates gene transcription through recognition of RA response elements (RARE) (*21–23*).

RAREs are dispersed throughout the genome on promoters, enhancers and intragenic regions and regulate thousands of genes that result in pleiotropic biological effects ranging from neuronal development to bone morphology (*20, 24, 25*). Vitamin A and RA are also very important for the development and function of our immune system (*17*). Thus, RA promotes the formation of SLOs (*26, 27*) during embryogenesis, it induces expression of the gut homing molecules CCR9 and α4β7 on T cells and innate lymphoid cells (*28–30*) and is critical for the differentiation of T-helper (T_H_) cells and the balance between T_H_ cell subsets (*31–35*). Furthermore, RA regulates T_H_1 to T_H_17 plasticity (*36*) and is critical for the development of small intestinal intra-epithelial lymphocytes (*37, 38*), while more recent data showed the importance of RA in establishing tissue resident memory cells (*39, 40*). Although treatment of cells or mice with synthetic RA can regulate some functions of γδ T cells (*41, 42*), the intrinsic requirements for RA in this cell population has not been elucidated.

Herein we provide evidence that RA is a key molecular determinant throughout the life of γδT17 cells. We show that during embryonic life, RA strengthens the TCR signal to limit γδT17 generation; during neonatal life RA is important for survival, while during adult steady-state conditions it maintains γδT17 numbers in SLOs. In the context of central nervous system inflammation, modelled by autoimmune experimental encephalomyelitis (EAE), RA is necessary to sustain γδT17 cells and their ability to produce cytokines, as well as to accumulate in the brain.

Through combined defects in the T_H_17 compartment, mice with impaired RAR activity in type 3 lymphocytes show near complete resistance to EAE symptoms.

## Results

### RA limits the development of γδT17 cells

In order to test the intrinsic importance of RA in γδT17 cells we crossed RORγt-Cre recombinase^+^ mice (RORγt^CRE^) (*43*) with mice whereby constitutive expression of a RARα dominant negative (dn) transgene within the *ROSA26* locus was prevented by a floxed stop codon (RARdn^F/F^) (*44*).

Excision of the stop codon allowed constitutive RARdn transgene expression, preventing endogenous RARα from becoming activated (*44*) (**Fig. 1A**). Using RORγt^CRE^-RARdn^F/F^ mice (referred to thereafter as Cre^+^) and their Cre^−^ littermate controls we first assessed whether RA is implicated in thymic development. Embryonic thymic γδT17 cells were identified as CD24^−^CD27^−^CD45RB^−^CD44^+^ (*6*), and in the neonates as CD27^−^CD44^+^ (*4*) and were split into Vγ4^+^ and Vγ4^−^ (Vγ nomenclature according to Heilig & Tonegawa (*45*)) (**Fig. S1A**). At embryonic day e18 total γδ T cells were not affected by inactive RARα (**Fig. S1B**), however, we found that numbers of γδT17 cells, of which approximately 90% were Vγ1/4^−^ (**Fig. S1C**), were elevated, albeit not significantly (**Fig. S1D**). At that point, very few Vγ4^+^ cells could be detected and their numbers were unchanged (**Fig. S1D**). One day after birth, throughout neonatal life and until weaning at 3 weeks, the numbers of Vγ4^−^ γδT17 cells were increased in Cre^+^ thymi (**Fig. S1E**). Of note, while the numbers of Cre^−^ Vγ4^−^ γδT17 cells plummeted shortly after birth, Cre^+^ cells persisted in the thymus for 2-3 weeks (**Fig. S1E**). There were no observable changes in Cre^+^ Vγ4^+^ cells up until day 21 after birth, at which point their numbers indicated a small decline (**Fig. S1E**).

**Figure 1.**
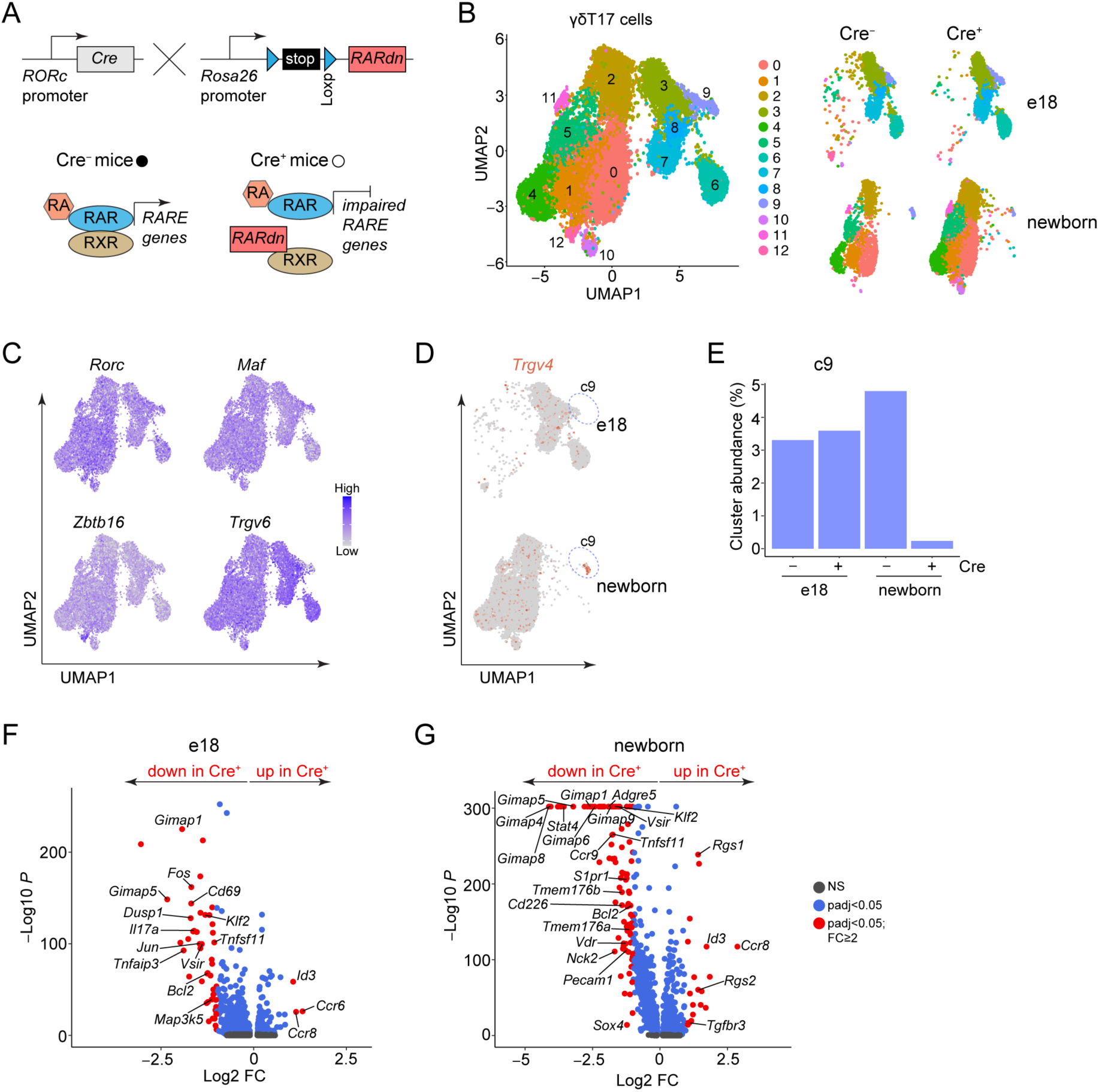
**RA sustains multiple transcriptional programs during γδT17 cell development.** (**A**) Schematic representation of the mouse model. RORγt^CRE^-mediated expression of the RARdn transgene prevents RARα from interacting with RXR and thus cannot initiate its transcriptional activity. (**B**) UMAP plot displaying integrated e18 and newborn γδT17 thymocytes from transgenic (Cre^+^) and littermate control (Cre^−^) mice after unsupervised clustering (left) and UMAP split based on genotype and timepoint (right). (**C**) Feature plots illustrating the distribution and expression levels of *Rorc*, *Maf*, *Zbtb16* and *Trgv6*. (**D**) Feature plot for *Trgv4* expression split based on timepoint with cluster c9 circled. (**E**) Proportions of cluster c9 within each timepoint and genotype, shown as a fraction of the total γδT17 cell population. (**F-G**) Volcano plots for DEGs determined using a Wilcoxon Rank Sum test between Cre^+^ and Cre^−^ cells for e18 (**F**) and newborn cells (**G**). NS: not significant; FC: fold change; log_2_FC = 1, representing a fold change of at least 2. Selected DEGs are labeled on the plot.

The increased numbers of Vγ4^−^ γδT17 cells both in embryonic and neonatal thymus suggested that RA may negatively regulate the development of these cells. In order to start gaining mechanistic insights that explained this phenotype, we set up fetal thymic organ cultures (FTOCs) as previously described (*6*), in the presence or absence of a synthetic RARα inhibitor (**Fig. S1F**). We found that RARα inhibition promoted the generation of higher numbers of γδT17 cells (**Fig. S1G**), which under these conditions are dominated by the Vγ4^−^ subset (*6*). However, exogenous high concentration of RA in the culture with all-trans RA (ATRA) did not reduce γδT17 cell numbers (**Fig. S1H**). Collectively, these data show that impaired sensing of RA favors thymic output of γδT17 cells and suggest that sustained RARα activity may limit their embryonic development.

### RA sustains multiple transcriptional programs during γδT17 cell development

The above data prompted us to further investigate how RA may influence the biology of γδT17 cells during embryonic and early life development. To this end, we isolated γδT17 cells from e18 and newborn Cre^−^ or Cre^+^ thymi (see gating strategy in **Fig. S1A**) and performed single-cell RNA- sequencing experiments (scRNA-seq). We used Uniform Manifold Approximation and Projection (UMAP) analysis to visualize γδT17 cells by integrating both time points and genotypes.

Unsupervised clustering showed 13 clusters, 5 unique to e18 (c3, c6, c7, c8, c12), 7 unique to newborn (c0, c1, c2, c4, c5, c10, c11) and one cluster (c9) that was common (**Fig. 1B and Table S1**). Expression of signature γδT17 genes such as *Rorc*, *Zbtb16* and *Maf* was found in all clusters, while the majority of cells expressed *Trgv6* in line with Vγ6^+^ cells being more abundant in the thymus at these stages of life (**Fig. 1C**). *Trgv4* expressing cells were found mostly in cluster c9, which in newborn mice was almost entirely derived from Cre^−^ cells (**Fig. 1D-E**). Although this suggested that RA supported the early development of Vγ4^+^ cells, their numbers were not affected in either e18 or newborn Cre^+^ thymus (**Fig. S1D-E**).

Next, we investigated differentially expressed genes (DEGs) for both e18 and newborn thymus between Cre^−^ and Cre^+^ cells. At e18 there were 43 genes downregulated and 3 upregulated in Cre^+^ cells, while in the newborn there were 90 genes downregulated and 20 upregulated in Cre^+^ cells (**Fig. 1F-G and Table S2**). Some of the downregulated genes at e18 but not newborns included the AP-1 transcription factors *Jun* and *Fos* and the signaling molecules *Dusp1*, *Map3k5* and *Tnfaip3*, as well as *Il17a* and *Cd69* (**Fig. 1F-G**). Both embryonic and newborn thymic Cre^+^ cells had reduced levels of *Tnfsf11* (RANKL) and *Vsir* (VISTA) while newborn Cre^+^ cells had reduced *Adgre5* (CD97), *Cd226*, *Vdr*, and *Stat4* (**Fig. 1F-G**), suggesting a more generalized defect in T cell function. The gene coding the transcription factor KLF2 was also downregulated in both embryonic and newborn Cre^+^ cells, while its target *S1pr1*, which promotes thymic egress (*46*), was additionally decreased in newborn Cre^+^ thymocytes (**Fig. 1F-G**). Notably, Cre^+^ cells had downregulated many genes of the GIMAP family (e.g. *Gimap1*, *Gimap4*, *Gimap5*), especially in the newborn (**Fig. 1F-G**). GIMAPs are GTPases implicated in the development and differentiation of T cells and have potent pro-survival and anti-apoptotic roles (*47*). In line with this, the gene coding the anti-apoptotic molecule Bcl-2 was also downregulated in the absence of active

RARα in the embryo and newborn (**Fig. 1F-G**). Collectively, in the embryonic and newborn thymus RA regulates a transcriptional program in γδT17 cells related to multiple signaling pathways, tissue homing and retention as well as cell survival.

### RA strengthens the TCR signal during embryonic γδT17 cell development

Next, we wanted to understand how can the above transcriptional changes explain the increased thymic output of γδT17 cells in mice with inactive RARα. We noted that the genes *Jun*, *Fos*, *Dusp1* and *Map3k5*, which have all been implicated in TCR signaling, were specifically downregulated in Cre^+^ e18 cells (**Fig. 1F-G**). We additionally found that cluster c12 separated from the rest based on 1351 DEGs, a large proportion of which appeared to be associated with cellular signaling, such as *Cblb*, *Malt1*, *Fyn*, and *Traf3* (**Table S1**). We then used over-representation analysis (ORA) to test whether the genes in cluster c12 enriched specific signaling cascades of immunological importance, and found that the TCR signaling pathway was the most highly and significantly enriched (**Fig. 2A**). Furthermore, cluster c12 was unique to e18 thymocytes and it was almost absent in Cre^+^ cells (**Fig. 2B and Fig. 1B**). Thus, these data indicate that RA is important in the embryonic thymus to induce genes that are corelated with TCR induced pathways.

**Figure 2.**
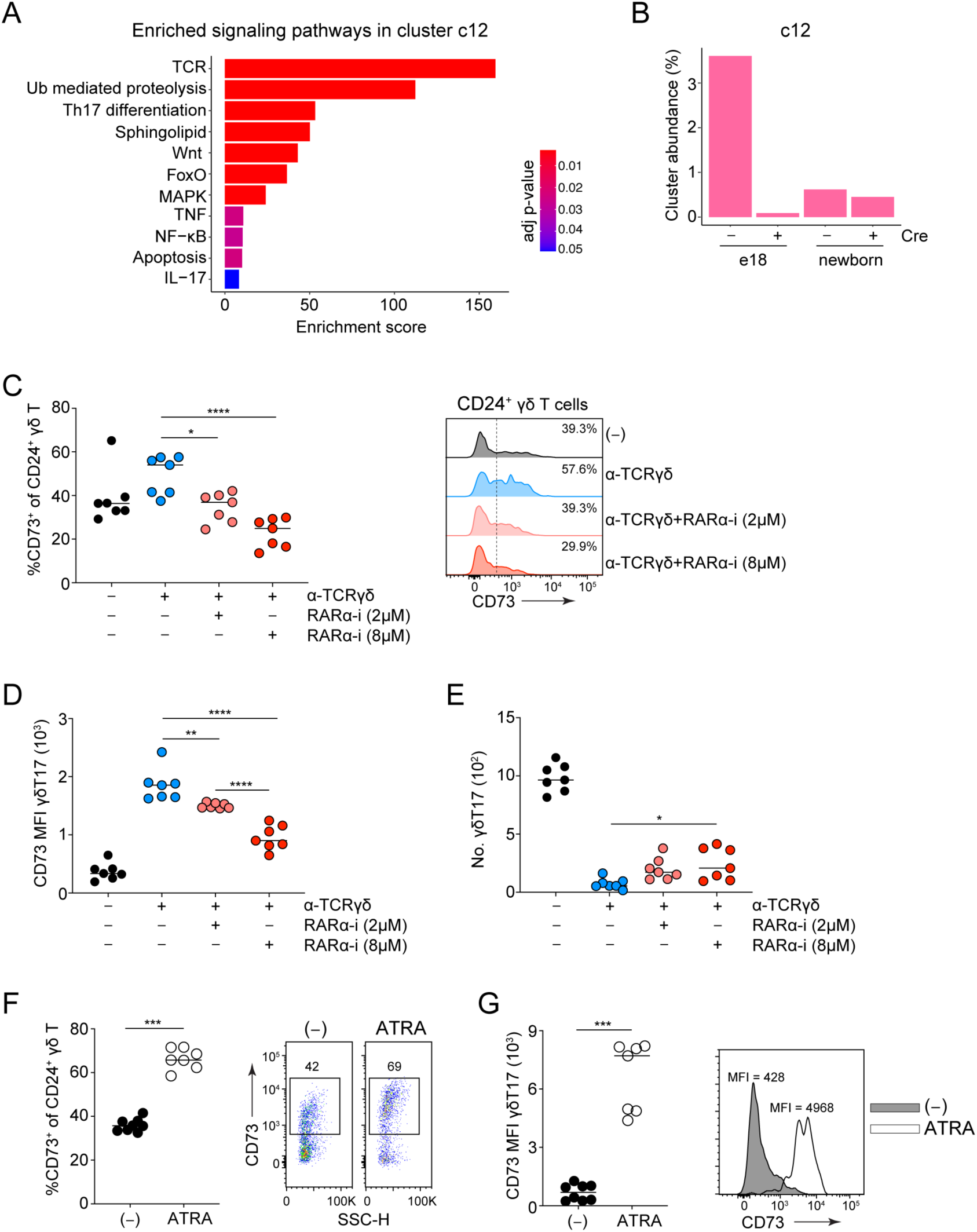
**RA strengthens the TCR signal during embryonic γδT17 cell development.** (**A**) KEGG pathway enrichment analysis of DEGs in cluster c12 compared to all other clusters performed using enrichR. Pathways are color-coded by adjusted p-value, with red indicating greater statistical significance. The combined enrichment score is shown on the x-axis. (**B**) Proportions of cluster c12 within each timepoint and genotype shown as a fraction of the total γδT17 cell population. (**C-E**) FTOCs were set up as described in **Fig. S1F** and were treated with α- TCRδ in the presence or absence of 2 and 8 μM RARα inhibitor (RARα-i). Frequencies of CD73^+^ cells within the immature (CD24^+^) γδ T cell population across the different treatment combinations (left), and representative ridgeline plots of CD73 staining (right) (**C**). MFI of CD73 staining in the γδT17 cell population (**D**). Numbers of γδT17 cells in the FTOCs across the treatments (**E**). (**F-G**) In separate experiments, FTOCs were treated with 160 nM of ATRA or left as untreated controls (−). Frequency of CD73^+^ cells in the CD24^+^ γδ T population (left) and representative FACS plots of CD73 staining of the same population (right) (**F**). MFI quantification of CD73 staining in γδT17 cells (left) and representative histogram of CD73 staining of the same population (right) (**G**). In (**C-G**) each symbol represents 2-3 thymic lobes pooled and the line the median; data are pool of two experiments; p-values were determined using One-Way ANOVA with Tukey’s multiple comparisons test for **C-E** and Mann–Whitney test for **F-G**; *P < 0.05. **P < 0.01; ***P < 0.001; ****P < 0.0001.

The above led us to hypothesize that RA through RARα may regulate TCR signaling. To test this hypothesis, we set up FTOCs in the presence of a RARα inhibitor and the agonistic anti(α)- TCRγδ GL3 Ab, which triggers the TCR and suppresses γδT17 development (*6, 48*). The surface nucleotidase CD73 can be used as a surrogate marker to indicate TCR activity in early γδ T cell committed progenitors (*49*). α-TCRγδ stimulation induced expression of CD73 in both immature CD24^+^ γδ T and committed γδT17 cells, which was antagonized by RARα inhibition (**Fig. 2C-D**), and under these conditions, γδT17 output was partially rescued (**Fig. 2E**). In contrast, addition of ATRA increased surface expression of CD73 in both CD24^+^ γδ T (**Fig. 2F**) and γδT17 cells (**Fig. 2G**). Collectively, these data show that impaired RA sensing attenuates TCR signaling in embryonic life and help explain the increased numbers of γδT17 in Cre^+^ thymi. Our findings suggest that under normal conditions RA strengthens TCR signaling in the embryonic thymus, limiting thus the development of γδT17 cells.

### RA promotes survival and represses apoptosis of neonatal γδT17 cells

Attenuated TCR signaling helps explain the increased numbers of γδT17 cells in the embryonic and neonatal thymus. However, higher thymic output did not translate into higher numbers in the periphery. In contrast, Vγ4^−^ γδT17 numbers were not increased in the neonatal lymph nodes (LNs), while the numbers of Vγ4^+^ γδT17 cells were significantly reduced at 2 and 3 weeks after birth (**Fig. S2A**). Furthermore, adult Cre^+^ LNs contained normal numbers of Vγ4^−^ and reduced numbers of Vγ4^+^ cells (**Fig. S2B**), and in the adult spleen the numbers of both subsets were significantly reduced (**Fig. S2C**). We therefore hypothesized that there must be a compensatory mechanism by which the numbers of overtly generated γδT17 cells return to baseline or below. As we saw GIMAPs and *Bcl2* being downregulated in newborn Cre^+^ cells, we speculated that RA additionally supports survival and that when its sensing is inefficient, cells die at a higher rate under homeostatic conditions.

We initially examined in detail the DEGs (see **Table S2 and Fig. 1F-G**) between Cre^−^ and Cre^+^ cells at e18 and newborn thymus in order to pinpoint genes that have been reported before in the literature to have anti- or pro-apoptotic properties (**Table S3**). We identified 24 genes with anti- and 7 genes with pro-apoptotic properties and 3 with both (**Fig. 3A**). The majority of anti-apoptotic genes were differentially and highly expressed in Cre^−^ cells of newborn origin, although this was also true for the pro-apoptotic genes (**Fig. 3A**), indicating that RA is impacting the survival of γδT17 cells. As Bcl-2 is a key pro-survival and anti-apoptotic molecule, we then determined its protein levels in the thymus and LNs of newborn and neonatal mice. We found that Bcl-2 levels were significantly reduced in Vγ4^−^ and Vγ4^+^ γδT17 cells that could not sense RA (**Fig. 3B**). To investigate whether reduced levels of Bcl-2 were related to weaker TCR signals, we cultured newborn thymocytes with α-CD3 in an attempt to restore the TCR signal strength. We found that α- CD3 treatment induced Bcl-2 levels in Cre^+^ cells (**Fig. 3C**), suggesting that weak TCR signaling may contribute to impaired survival. To formally test whether RARα inactivity resulted in increased cell death, we analyzed the expression of the execution caspases 3 and 7. In order to induce measurable levels of apoptosis we stressed newborn thymocytes by culturing them in vitro for 40 hours in the absence of any survival factors. Under these conditions, we found a higher proportion of caspase 3/7 positive Cre^+^ γδT17 cells (**Fig. 3D**). Collectively, our data show that RARα sustains the survival of newly generated γδT17 cells and indicate that this is linked to its ability to regulate the strength of the TCR.

**Figure 3.**
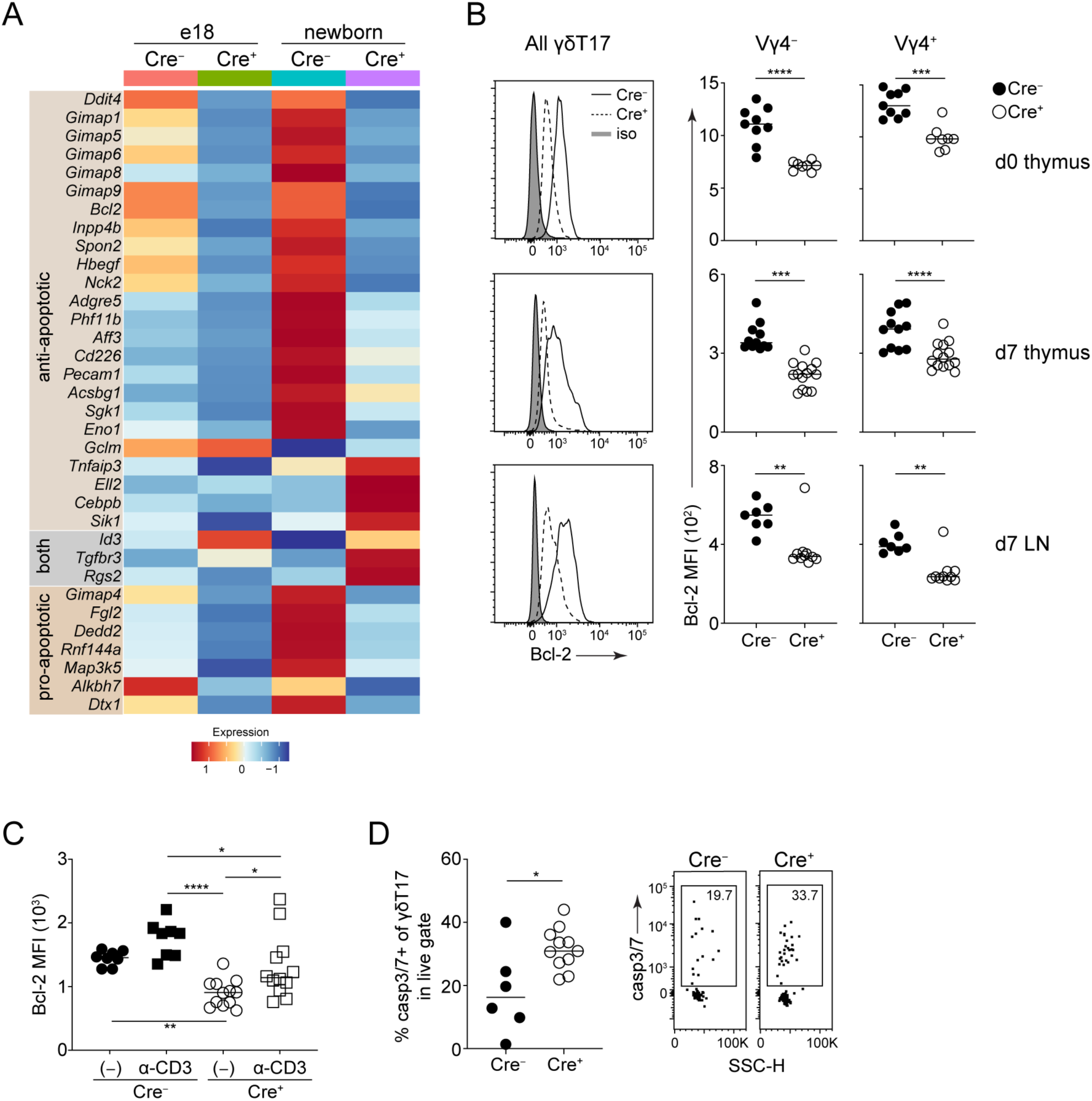
**RA promotes survival and represses apoptosis of neonatal γδT17 cells.** (**A**) Heatmap representation of selected DEGs between Cre^−^ and Cre^+^ cells for each timepoint grouped as anti-apoptotic, pro-apoptotic or both (**Table S3**). Colors indicate scaled gene expression levels, with red representing higher expression. (**B**) Quantification of Bcl-2 by flow cytometry for newborn (d0) thymus and neonatal (d7) thymus and lymph node (LN). Data is shown as representative histograms for γδT17 cells (left) and quantified individually for Vγ4^−^ and Vγ4^+^ cells within the γδT17 population. (**C**) MFI of Bcl-2 after overnight culture of newborn Cre^−^ and Cre^+^ thymocytes in the absence (−) or presence of 1 μg/ml α-CD3. (**D**) Frequency of caspase 3/7^+^ cells among the live γδT17 population following 40 hours of culture, along with a representative FACS plot (right). In (**B-D**) each symbol represents one mouse and the line the median; data are pool of three experiments; p-values were determined using One-Way ANOVA with Tukey’s multiple comparisons test for **C** and Mann–Whitney test for **B** and **D**. *P < 0.05; **P < 0.01; ***P < 0.001; ****P < 0.0001.

### RA induces activation, survival and interferon gene signatures in inflammation

Next, we wanted to investigate whether RA is also required in γδT17 cells during inflammatory responses. As γδT17 cells, in particular Vγ4^+^, are important for the development and progression of MOG-induced EAE (*50*), we isolated γδT17 cells from the spleens of Cre^−^ and Cre^+^ mice that were either unimmunized (day 0, d0) or had been immunized 11 days prior (d11), performed scRNA-seq, and analyzed Vγ4^+^ and Vγ6^+^ cells separately. By integrating both time points and genotypes, we identified 7 Vγ4^+^ clusters (**Fig. 4A**), all of which expressed type 3 genes (e.g. *Rorc*, *Maf*, *Il23r*), and none expressed *Cd27*, a marker of type 1 cells (**Fig. 4B and Table S4**). All clusters expressed *Trdv2-2*, except cluster c3 which was *Trdv5*^+^ and had the lowest levels of *Il17a* (**Fig. 4B and Table S4**), while cluster c6 was contaminated with αβ T cells (**Fig. 4B and Table S4**). Clusters c2 and c4 had distinct proliferative signatures (**Fig. 4B and Table S4**) and showed increased abundance in EAE, however, considerably more in Cre^+^ mice (**Fig. 4C**). Interestingly, cluster c1 expressed a number of IFN-induced genes (e.g. *Nkg7*, *Irf7*, *Gbp6*, *Zbp1*) together with *Tbx21*, downregulated *Ccr6* and was the cluster whose abundance increased after EAE and only in Cre^−^ mice (**Fig. 4B-D**). Additionally, cluster c1 had detectable *Ifng* and reduced levels of *Il17a* compared to other clusters (**Fig. 4B, D**).

**Figure 4.**
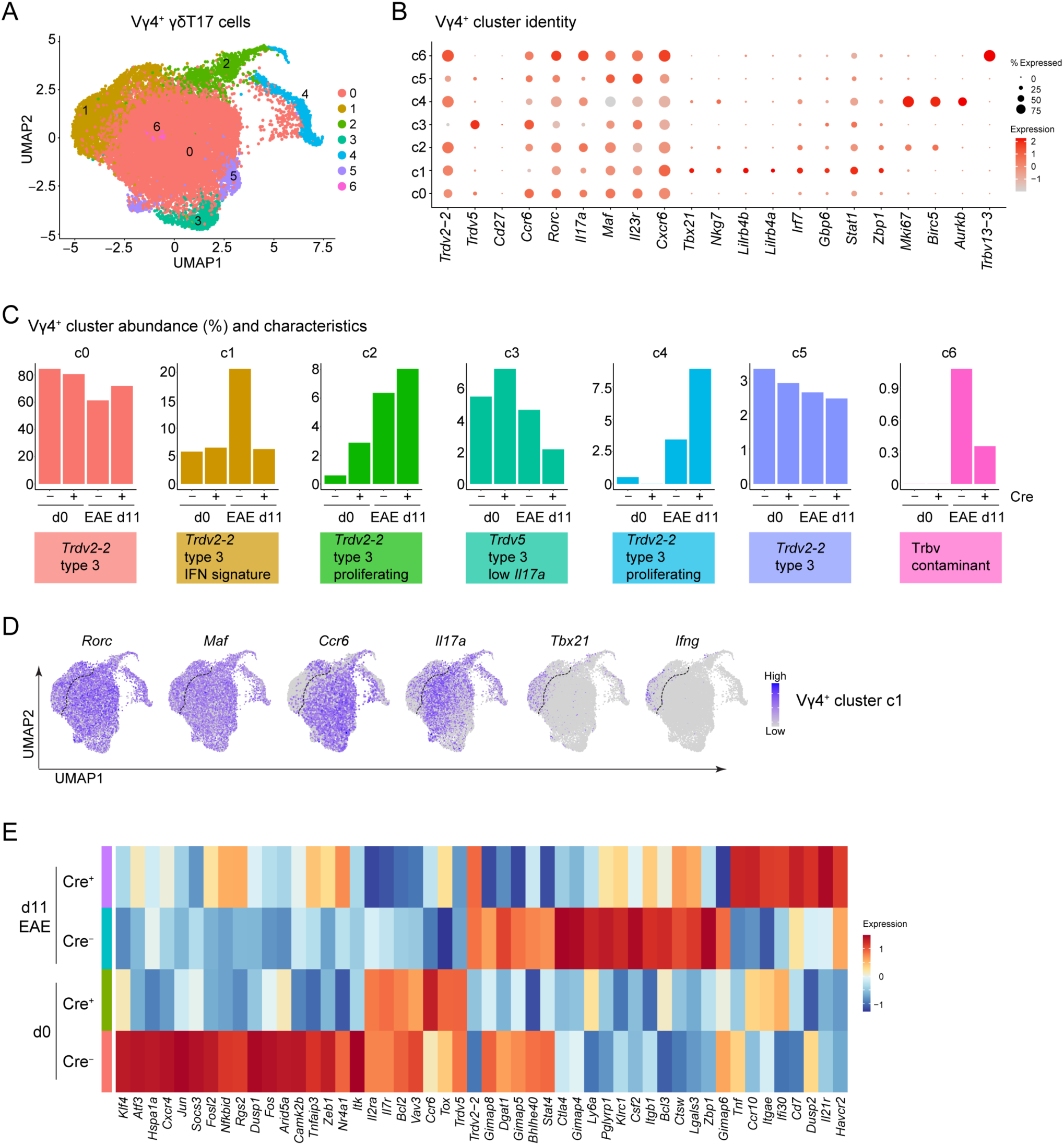
**RA induces activation, survival and interferon gene signatures in inflammation.** (**A**) UMAP of Vγ4^+^ γδT17 clusters integrated from the spleen of Cre^−^ and Cre^+^ mice before (d0) and 11 days (d11) after MOG_35-55_ immunization. (**B**) Dot plot showing the scaled expression of selected genes across clusters, with red representing higher expression. Dot size indicates the percentage of cells within each cluster expressing the corresponding gene. (**C**) Cluster characterization and abundance within each genotype and timepoint, shown as a fraction of all Vγ4^+^ cells. (**D**) Feature plots illustrating the distribution and expression levels of *Rorc*, *Maf*, *Ccr6*, *Il17a*, *Tbx21* and *Ifng*. A line is drawn to delineate the IFN signature cluster c1 from the others. (**E**) Heatmap representation of selected DEGs between Cre^−^ and Cre^+^ cells for each EAE timepoint. Colors indicate scaled gene expression levels, with red representing higher expression.

Analysis of DEGs showed that at d0, Cre^+^ cells had a clear reduction in genes involved in TCR, cytokine and NF-κB signaling (e.g. *Jun*, *Fos*, *Nr4a1*, *Tnfaip3*, *Itk*) (**Fig. 4E and Table S5**), suggesting that when RARα is inactive Vγ4^+^ γδT17 cells have impaired propensity to become activated. As such, at d11, it was mostly Cre^−^ cells that upregulated genes classically associated with activated T cells and IFN signaling such as *Ctla4*, *Ly6a*, *Stat4*, and *Zbp1*, in addition to the cytokine *Csf2* and the pro-survival GIMAP genes *Gimap5*, *Gimap6*, and *Gimap8* together with *Bcl3* (**Fig. 4E and Table S5**). In contrast, genes that discerned Cre^+^ cells responding to immunization included the adhesion molecules *Ccr10* and *Itgae*, *Il21r*, and *Tnf* (**Fig. 4E and Table S5**). We did not find any proliferation or cell cycle related genes that were downregulated in Cre^+^ cells suggesting intact proliferative capacity (**Table S5**). Collectively, these data show that RARα maintains a status of activation and survival without interfering with cell proliferation, and that during EAE supports Vγ4^+^Vδ2-2^+^ γδT17 cells characterized by an IFN gene signature.

When we investigated Vγ6^+^ cells, we identified 6 clusters, all with a type 3 identity, of which c4 was a *Scart2*^+^ contaminant (**Fig. S3A-B and Table S4**). Cluster c1 expressed higher levels of *Egr1*, and was nearly absent in Cre^+^ cells prior to immunization (**Fig. S3B-C and Table S4**).

Cluster c2 could be distinguished by high levels of *Tnfsf11* and *Stab2* as well as lower *Il17a*, and both before and after immunization was more prevalent in Cre^+^ cells (**Fig. S3B-C and Table S4**). Cluster c3, similar to Vγ4^+^ cluster c1, downregulated *Ccr6*, expressed *Tbx21* and had a distinct IFN signature (**Fig. S3B and Table S4**). As was the case for the Vγ4^+^ counterpart, this cluster had reduced levels of *Il17a* and detectable transcripts of *Ifng* and although its abundance did not increase in EAE, it was more prevalent in Cre^−^ cells (**Fig. S3B-D**). Cluster c5 was of very low frequency, had a proliferation related gene signature and was more abundant at d11 after EAE induction in Cre^+^ mice (**Fig. S3C**). Cre^−^ Vγ6^+^ cells appeared to be relatively unresponsive to EAE, with the exception of a modest increase observed for cluster c0 (**Fig. S3C**), while the abundance of cluster c1 was much higher in Cre^+^ cells after immunization (**Fig. S3C**).

Analysis of DEGs among genotypes and time points revealed transcriptional changes relatively similar to Vγ4^+^ cells (**Table S5**). Thus, at d0, Cre^−^ cells had high expression of many genes related to immune signaling pathways (e.g. *Nr4a1*, *Dusp2*, *Rel*), immune cell activation (e.g. *Il17a*, *Cd69*, *Itga4*) and survival (*Gimap8*, *Gimap9*, *Bcl2a1d*), indicating a better cellular fitness during immune responses (**Fig. S3E and Table S5**). A lot of these genes were downregulated at d11 of EAE and a fraction was now more highly expressed in Cre^+^ cells (**Fig. S3E and Table S5**). Instead, Cre^−^ Vγ6^+^ cells showed induction of IFN-associated genes after immunization (e.g. *Ly6a*, *Ifi47*, *Stat4*) (**Fig. S3E and Table S5**). These data suggest that Vγ6^+^ cells with active RARα are better equipped to survive and react to inflammatory stimuli, they too show an IFN gene signature, however, they do not respond well to EAE.

### RA is necessary for EAE mediated neuroinflammation

The above data suggested that γδT17 cells with inactive RARα would be defective in mounting a response during EAE. To test this, we immunized mice and analyzed cell numbers and cytokine production in the spleen. We found that at d11 post immunization, there was a dramatic reduction in the numbers of Cre^+^ Vγ4^+^ cells (**Fig. 5A**). While the number of normal Vγ4^+^ γδT17 cells increased on average 19-fold by d11, their Cre^+^ counterparts increased 9-fold (**Fig. 5A**). At the same time, there was a profound loss of IL-17A-producing cells (**Fig. 5B and Fig. S4A**). To correct for the highly reduced numbers of Cre^+^ cells, we assessed the frequency of IL-17A^+^ cells within the Vγ4^+^ γδT17 population. We found that Cre^+^ Vγ4^+^ γδT17 cells were compromised in their ability to produce IL-17A (**Fig. 5C**). The EAE-induced increase of Vγ4^−^ γδT17 cell numbers was considerably less prominent, and the impact of RARα was insignificant (**Fig. S4B**), which agreed with our scRNA-seq data (**Fig. S3C**). We additionally observed that Vγ4^+^ cells could produce low amounts of IFN-γ together with IL-17A and this was RA-dependent (**Fig. 5D-E**). This may represent the type 1 *Ifng*-expressing c1 cluster in our scRNA-seq dataset (**Fig. 4C-D**). Although we had observed a similar cluster (c3) in Vγ4^−^ cells (**Fig. S3C-D**), we could not detect IFN-γ protein by flow cytometry (**Fig. S4C-D**). By d21 post immunization, the γδT17 response had subsided (**Fig. 5A** and **Fig. S4B**). Concomitantly, we found that Cre^+^ mice developed significantly reduced paralysis symptoms (**Fig. 5F**) and retained their body weight (**Fig. 5G**). This profound resilience to developing EAE disease was puzzling, since mice deficient in all γδ T cells have delayed disease but still develop evident symptoms (*51*). We thus hypothesized that the phenotype of RORγt^CRE^- RARdn^F/F^ mice could be explained by an additional defect in T_H_17 cells, which are necessary for EAE pathogenesis (*52*), and which get help from γδT17 cells (*50, 51*). In this regard, we found defects in the CD4^+^ T cell compartment at d11 of EAE, coinciding with the defective γδT17 response. Hence, although numbers of activated CD44^+^CD4^+^ T cells were only slightly reduced at d11 (**Fig. S4E**), there was a significant impairment in their ability to produce IL-17A and IFN-γ (**Fig. S4F-I**). Thus, RA is important for the full development of EAE-associated symptoms and for Vγ4^+^ γδT17 and T_H_17 cells to efficiently respond to neuroinflammatory stimuli.

**Figure 5.**
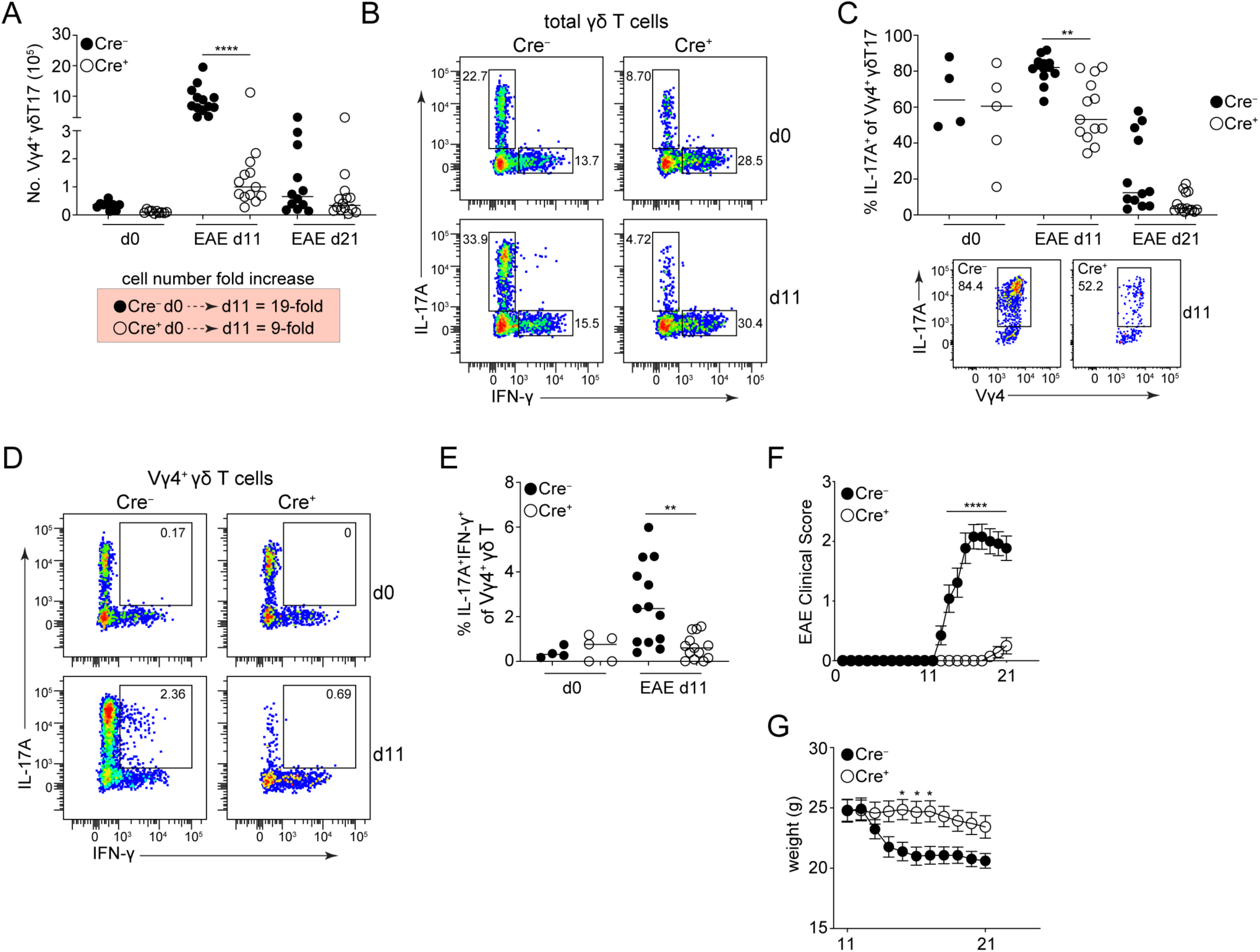
**RA is necessary for EAE mediated neuroinflammation.** (**A**) Numbers of Vγ4^+^ γδT17 cells in the spleen at d0, d11 and d21 after MOG_35-55_ immunization. The fold increase in cell numbers from d0 to d11 for each genotype is also shown. (**B**) Representative FACS plot showing IL-17A and IFN-γ cytokine production in the spleen within the total γδ T cell population at d0 and d11. (**C**) Frequency of IL-17A^+^Vγ4^+^ cells within the γδT17 population at different timepoints of EAE. Representative FACS plot for the IL-17A staining is also shown. (**D**) Representative FACS plots for IL-17A and IFN-γ co-production in the spleen by Vγ4^+^ γδ T cells. (**E**) Frequency of IL-17A^+^IFN-γ^+^ cells within the Vγ4^+^ γδ T cell population. (**F**) Disease progression measured by clinical symptoms until day 21 post immunization. (**G**) Mouse weight between d11 and d21. In **F** and **G** data are a pool of 13 (Cre^−^) or 14 (Cre^+^) mice and shown as mean±sem. Statistical analysis was performed using 2-way ANOVA with Bonferroni’s multiple comparisons test. For all the other plots, each symbol represents a mouse and the line the median; data are a pool of three experiments; p-values were determined using One Way ANOVA with Tukey’s multiple comparisons test. *P < 0.05; **P < 0.01; ****P < 0.0001.

### RA is required for α4β1 expression and γδT17 cell infiltration to the brain

A key role of RA in the immune system is the imprinting of the gut homing molecules CCR9 and α4β7 on T cells (*17*). We had also observed that RA regulated genes related to cell adhesion and migration in thymic γδT17 cells (**Fig. 1 and Table S2**). Thus, we examined the DEGs between Cre^+^ and Cre^−^ Vγ4^+^ γδT17 cells at d11 of EAE and identified transcripts related to cell adhesion, migration or tissue retention (**Table S6**). We identified 19 such genes, 4 of which were higher and 8 were lower in Cre^+^ cells at d11, while 7 remained unchanged (**Fig. 6A**). Differences included the downregulation of *Rasa3*, important for T cell entry and exit from SLOs but also T cell responses and T_H_17 pathology during EAE (*53, 54*), and the upregulation of *Rgs1*, which negatively regulates chemokine receptor signaling (*55*) (**Fig. 6A**). Notably, *Itgb1*, which codes the β1 integrin chain, was also significantly downregulated in Cre^+^ cells (**Fig. 6A**). The β1 can pair with the α4 chain to form the α4β1 integrin, also known as VLA-4 (*56, 57*), which is important for the homing of pathogenic lymphocytes to the brain during EAE and has been used as a therapeutic target in multiple sclerosis (MS) (*58–60*). We therefore wanted to test whether levels of α4β1 are altered in cells with inactive RARα. To do this, we co-stained with antibodies that mark the α4β7 complex, and either the α4 or β1 chains at d0 and d11 of EAE. Hence, by gating out α4β7^+^ cells and focusing on cells co-expressing the α4 and β1 chains, we could discern α4β1^+^ cells. We found that γδT17 cells expressed significantly less α4β1 when RARα was inactive (**Fig. 6B-C**), and of the α4β1- expressing γδT17 population over 80% were Vγ4^+^ (**Fig. 6D**). Reduced expression of α4β1 was also evident in CD44^+^CD4^+^ T cells (**Fig. S5A-B**). Failed induction of surface α4β1 correlated with impaired γδT17 and CD44^+^CD4^+^ T cell infiltration into the brain (**Fig. 6E and Fig. S5C**). Although the low cell numbers in the brain might simply reflect the low numbers of responding cells in the periphery, it helps to further explain the profound resistance of RORγt^CRE^-RARdn^F/F^ mice to EAE.

**Figure 6.**
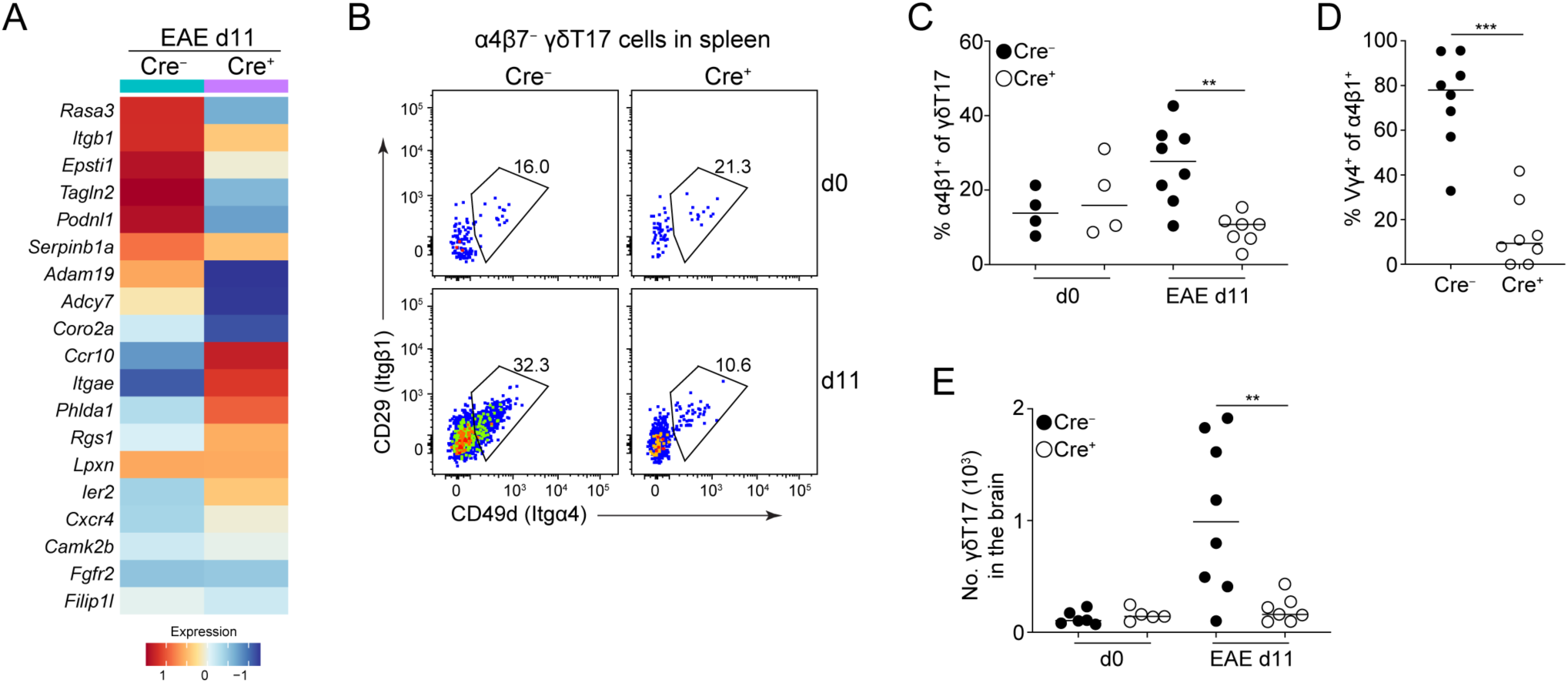
**RA is required for α4β1 expression and γδT17 cell infiltration to the brain.** (**A**) Heatmap representation of selected DEGs with potential roles in migration (**Table S6**) between Cre^−^ and Cre^+^ Vγ4^+^ cells at d11. Colors indicate scaled gene expression levels, with red representing higher expression. (**B**) Representative FACS plots of α4β1 integrin expression on splenic γδT17 cells at d0 and d11 of EAE (cells were pre-gated as α4β7^−^). (**C**) Frequency of α4β1^+^ cells within the γδT17 population. (**D**) Frequency of Vγ4^+^ cells within the α4β1^+^ γδT17 population at d11 of EAE. (**E**) Numbers of γδT17 cells in the brain of Cre^−^ and Cre^+^ at d0 and d11 of EAE. In all graphs, each symbol represents one mouse; data are pool of two experiments; p-values were determined using One-Way ANOVA with Tukey’s multiple comparisons test for **C** and **E**, and Mann–Whitney test for **D**. *P < 0.05; **P < 0.01; ***P < 0.001.

## Discussion

In the present study we demonstrate that RA and its nuclear receptor RARα act on multiple levels to regulate the development, homeostasis and inflammatory responses of IL-17-producing γδ T cells. We found that cells unable to sense RA through RARα in the embryonic and newborn thymus undergo a transcriptional reorganization affecting a multitude of signaling modalities, homing and tissue retention potential in addition to survival. As such, our data suggest that RA strengthens TCR signaling in the embryonic thymus putting thus a break on γδT17 development, while after birth RA is critical to maintain survival of newly developed cells by counteracting apoptosis. In adult mice and during the course of EAE neuroinflammation, RA sustains a transcriptional program that indicates high propensity to respond to inflammatory stimuli and immunization, especially in Vγ4^+^ γδT17 cells. Thus, Vγ4^+^ γδT17 cells with inactive RARα failed to sufficiently increase in numbers or produce cytokines in response to EAE. This was accompanied by reduced expression of the integrin α4β1 and accumulation of cells in the brain. Consequently, and combined with a defective effector CD4^+^ T cell compartment, RORγt^CRE^-RARdn^F/F^ mice were almost resistant to EAE associated symptoms.

It is well-established that the strength of the TCR during thymic development is an important determinant for the lineage choices of γδ T cells. The current model suggests that cells who escape Skint-1 selection and do not become Vγ5^+^, receive either strong or weak TCR signals, the latter of which, through the PI3K-Akt pathway, determine their ability to become γδT17 (*5–7, 48*). In contrast, strong ERK-dependent TCR signals promote the generation of IFN-γ-producing γδT1 cells (*6*). Although Vγ4^+^ γδT17 appear less dependent on the TCR and are generated from pre- programmed Sox13^+^ DN1d progenitors, they too need TCR signaling to transition to fully mature IL-17-producing cells (*61*). Our data show that inactivation or inhibition of RARα reduces the TCR signaling strength, favoring thus the generation of γδT17 cells in the embryonic and newborn thymus. It is most likely that this is regulated at the gene expression level, since we found that cells with inactive RARα have impaired transcriptional modalities related to TCR, especially in embryonic life. Thus, gene expression and hence biological effects downstream of the TCR are impaired in the absence of RARα or RA sensing. Previous data in conventional T cells have also showed that RARα may cooperate with TCR mobilized transcription factors, such as NFATc2, to induce gene expression (*62*), suggesting a broader role of RA in fine tuning the outcome of TCR signaling. Recently, a novel RARα isoform located in the cytoplasm was shown to interact directly with the αβTCR and Zap70 to strengthen early signaling (*63*). Although there is no evidence that this isoform is relevant or expressed in γδ T cells, RARα inhibition in our FTOC experiments could have interfered with both nuclear and cytoplasmic RARα. This notwithstanding, RA and its interaction with RARα comprise a mechanism by which the strength of the γδTCR can be fine- tuned to balance lineage choice of developing γδ T cell subsets.

Besides TCR signaling, thymic γδ T cells that could not sense RA downregulated many genes related to T cell activation, adhesion, migration, tissue retention and survival. This was not surprising as RA and its nuclear receptors regulate thousands of genes in multiple cell types and tissues throughout life. Notably, there were many genes downregulated, especially in newborn γδT17 cells, related to survival, including *Bcl2* and a number of GIMAP transcripts, which led to increased levels of apoptosis. RA is a well-known regulator of cell death and survival, and although it has been shown to possess both pro- and anti-apoptotic activity, it is mostly considered as an inducer of apoptosis (*64*). However, RA can prevent apoptotic death in cardiomyocytes (*65*) as well as in developing brain cells (*44*), while early studies in T cells had suggested that RA could inhibit activation induced cell death (*66, 67*). Although the mechanisms by which RA may inhibit apoptosis in T cells have not been discerned, it is plausible that they are transcriptionally regulated through direct or indirect induction of genes that enhance survival and counteract cell death.

GIMAP genes for example have been shown to be transcriptional targets of RARα (*36*). It is also plausible that the survival defects that we observed in RARα inactive cells could be due to continuous weak TCR signaling, evident by the partial rescue of Bcl-2 in α-CD3 stimulated newborn γδT17 cells. In this regard, continuous weakening of the TCR signal will eventually result in the deletion of developing γδT17 cells (*48*). However, it is not yet known whether the γδTCR is actively signaling in the periphery and whether this is required to sustain homeostatic survival.

Collectively, the data suggest that through direct transcriptional targeting and by strengthening the γδTCR signal, RA and RARα impact in unison, and/or sequentially, thymic development and extrathymic survival of γδT17 cells.

T_H_17 and γδT17 cells are the major pathogenic populations in EAE (*51, 52, 68*) and their presence in MS has been correlated with disease severity (*69–72*). Hence, suppressing IL-17- secreting T cells could be beneficial in limiting MS pathology (*73–75*). Our data reveal that inability to sense RA reduces the propensity of γδT17 cells to become activated by regulating multiple transcriptional programs, including those associated with response to IFNs, survival, cytokine production and homing to the brain. The observation that IFN genes were readily inducible and RA-dependent, and that type 1-like γδT17 cells were highly abundant in EAE is reminiscent of T_H_ responses, which convert from type 3 to type 1 during disease progression (*76, 77*). Whether sensing of IFN or production of IFN-γ makes γδT17 subsets more pathogenic in EAE or it simply denotes activated cells has not been tested. Nevertheless, RORγt-Cre-mediated impairment of RA sensing led to a profound resistance in EAE disease. The data suggest that this was not only due to the aforementioned defects in γδT17 cells, but that parallel defects in T_H_17 cells contributed. In our transgenic model, RORγt^+^ type 3 cells develop in an “RA-less” environment, which we believe destabilizes them over time and become functionally incompetent. This raises the hypothesis that continuous exposure to low or high levels of dietary vitamin A from embryogenesis and neonatal to adult life may modulate fundamental transcriptional programs in type 3 cells impacting on their ability to mount immune responses. Thus, acutely altering the concentrations of available RA in competent cells may have very different outcomes. In this regard, administration of RA during ongoing EAE or pre-treatment of donor cells with RA prior to transfer into recipient mice suppressed the function of both T_H_17 and γδT17 cells and reduced disease severity (*41, 78*).

Consequently, despite the importance of RA in arming type 3 effector T cell populations during development and maturation, sudden exposure to high concentrations of RA may have the opposite effects.

Collectively, the data presented herein suggest that RA impacts γδ T cells from embryonic to neonatal to adult life and during inflammatory responses, and ensures their normal development, their survival and their ability to participate in immune responses. It would be of interest to explore whether low or high levels of dietary intake of vitamin A during pregnancy and infancy are able to modulate the biology of γδ T and other immune cells.

## Acknowledgments

We would like to thank everyone at the BioFacility at the Technical University of Denmark for continuous support with animal studies. The study was funded by the Lundbeck Foundation (R366- 2021-104) and the Novo Nordisk Foundation (NNF24OC0095974). NS and DP were funded by the BBSRC (BB/R017808/1).

## Author contributions

VB conceived the study, wrote the manuscript and analyzed data. MGFFdS and RA performed the experiments, analyzed the data and helped write the manuscript. JR, JF, ECF, MVB, DS and PPS helped with experiments, NS and DP performed all experiments related to FTOCs and helped write the manuscript.

## Materials and Methods

### Mice

All animal breeding and experiments were performed in-house at the BioFacility within the Technical University of Denmark and only after approval from the Danish Animal Experiments Inspectorate or at Queen Mary University of London in full compliance with UK Home Office guidelines. RORγt^CRE^ and *Rorc(γt)-Gfp^TG^* reporter (RORγt-GFP^+/−^) (*43*) mice were provided by Gerard Eberl. RARdn^F/F^ mice (*44*) were provided by William Agace after permission from San Sockanathan.

### Cell preparation and culture

Dissected adult, embryonic, newborn and neonatal lymphoid organs were isolated and crushed through a cell strainer and washed before being re-suspended in culture media (RPMI containing 10% FBS, Penicillin/Streptomycin, 0.1% β-ME, 20 mM HEPES and L-Glutamine) (ThermoFisher). An extra RBC lysis step was used for spleens. To isolate lymphocytes from the brain, mice were euthanized by CO_2_ asphyxiation, perfused with cold PBS and the whole brain was additionally rinsed in PBS. The tissue was cut in smaller pieces and digested for 30 minutes at 37 °C with 10 mg/ml DNase I (Sigma-Aldrich) and 10 mg/mL type IV collagenase from *Clostridium histolyticum* (Sigma-Aldrich), pipetting up and down every 10 minutes. Afterwards, the remaining tissue was homogenized with a Pasteur pipette and crushed through a cell strainer. Lymphocytes were separated using density gradient centrifugation with 37% Percoll (GE Healthcare) layered on 70% Percoll and centrifuged at 20 °C and 900 × *g* for 30 min with deceleration set to 0. Cells from the interphase were collected and washed once before used for staining. Cells from lymphoid organs were counted using a Sysmex XP-300. Cell counts from brain were determined using CountBright^TM^ Absolute Counting Beads (ThermoFisher).

### *Ex vivo* culturing of lymphocytes

For cytokine detection cells were cultured for 3.5 hours in the presence of 50 ng/ml PMA (Sigma- Aldrich, 750 ng/ml Ionomycin (Sigma-Aldrich) and 1 μl/ml BD GolgiStop (containing monensin). For determination of Bcl-2 levels after TCR stimulation, newborn thymocytes (isolated as described above) were cultured with or without 1 µg/ml soluble α-CD3 (clone 145-2C11, BioXCell) overnight (16 h). To measure the degree of apoptosis, the CellEvent Caspase-3/7 Green Flow Cytometry Assay Kit (Invitrogen) was used. In short, newborn thymocytes were cultured for 40h in culture media and 1 µl/ml of detection reagent was added 30 minutes before harvesting. In all cases, cells were incubated at 37°C, harvested and used for flow cytometry staining.

### Fetal thymic organ cultures (FTOCs)

Embryonic e15 thymic lobes from time-mated C57BL/6 or RORγt-GFP^+/−^ mice were cultured on nucleopore membrane filter discs (Whatman) in culture media (RPMI with 10% heat-inactivated FBS, Penicillin/Streptomycin, 0.1% β-ME, and L-Glutamine) (all reagents from ThermoFisher) for 7 days under control conditions or in the presence of either 160 nM ATRA (Sigma-Aldrich) or 2 or 8 µM of the selective RARα antagonist Ro 41-5253 (Sigma-Aldrich). In cultures containing Ro41, anti-TCRδ antibody GL3 (ThermoFisher) was added at a concentration of 0.5 μg/ml. All cultures were rested overnight in fresh media before analysis by flow cytometry.

### Experimental Autoimmune Encephalomyelitis

EAE was induced by sub-cutaneous injection of 50 μg of MOG_35-55_ peptide in CFA, while 200 ng pertussis toxin were injected intra-peritoneally (i.p.) on the day of immunization and 2 days later. From day 11 after immunization and until day 21, mice were weighed and scored for clinical signs as follows: 0: no symptoms; 1: tail paralysis; 1.5: impaired righting reflex; 2: paralysis of one hind limb; 2.5: paralysis of both hind limbs; 3: paralysis of one fore limb; 3.5: paralysis of one fore limb and weak second limb; 4: total limb paralysis. Mice were euthanized at day 21 after immunization. Lymphocytes from the brain and lymphoid organs were isolated as described above.

### Flow Cytometry

Cells were stained in U-bottom 96-well plates in 75 μl PBS containing 3% FBS with combinations of the following antibodies: CD4-FITC (RM4-4), CD19-FITC (6D5; Biolegend), CD8α-FITC (53-6.7), Vγ4-PerCP-eF710 (UC3-10A6; ThermoFisher), CD11b-PerCP-Cy5.5 (M1/70), IFNγ-APC (XMG1.2; BioLegend), CD29-APC (eBioHMb1-1; ThermoFisher), Vγ1-APC (2.11; Biolegend), CCR6-AF647 (140706), TCRβ-APCeF780 (H57-597; ThermoFisher), CD45RB-PE (16A), IFNγ-PE (XMG1.2; BioLegend), CD49d-PE (R1-2; ThermoFisher), Vγ4-PE (UC3-10A6; Biolegend); Bcl2-PE (BCL/10C4; Biolegend), α4β7-PECF594 (DATK32), CD3-PECF594 (145-2C11), CD24-PECy7 (M1/69), CD27-PECy7 (LG.3A10), CD73-PECy7 (BioTY/11.8; ThermoFisher), IFNγ-APC (XMG1.2), CD3-PECy7 (145-2C11), CD3-PE (145-2C11; BioLegend), RORγt-APC (B2D), TCRγδ- BV421 (GL3), CD44-V500 (IM7), CD45-V500 (30-F11), CD45.2-BV650 (104; Biolegend), CD27- BV650 (LG.3A10), IL-17A-BV786 (TC11-18H10), CD8α-BV786 (53-6.7), CD45-BUV395 (I3/2.3), CD19-Alexa Fluor 700 (1D3). Cells were stained for 30 minutes on ice and all antibodies were used at a 1:200 dilution. Prior to antibody staining, cells were incubated on ice with 100 µl of Fixable Viability Dye AF700 at a 1:1000 dilution in PBS. Cells were washed in 150 μl PBS containing 3% FBS in-between steps. Intracellular cytokine and transcription factor staining was performed using the eBioscience FoxP3 Transcription Factor Staining kit (ThermoFisher) according to the manufacturer’s instructions. Unless specified all antibodies and staining reagents were purchased from BD Biosciences. Samples were acquired on a BD LSR Fortessa using BD FACSDiva software v8.0.2. For sorting, the staining was carried out as described above but using PBS containing 0.5% BSA instead, and cells were sorted on BD FACSAria III.

### Single-cell RNA-seq

Thymic and splenic cells were prepared and stained as described above for cell sorting. However, cells were first incubated with 0.25 µg TruStain FcX PLUS (Biolegend) for 10 minutes on ice.

Without washing in between, cells from each mouse litter (3 for each timepoint) were hashed for 30 minutes on ice with TotalSeq-C0304, TotalSeq-C0305 and TotalSeq-C0306 antibodies (1 µg per 10^6^ cells) (Biolegend) respectively. For e18, only one litter was used. Afterwards, the antibody cocktail combined with the viability dye was directly added without a washing step and left for 30 minutes. Cells were then washed 3 times and sorted based on the gating strategy depicted in Figure S1A. Following the 10X Genomics Single Cell 5’ Chemistry (v2) platform, sorted γδT17 cells were loaded into a Chromium Controller and processed according to the manufacturer’s instructions. Library quality control and quantification were done using a 2100 Bioanalyzer equipped with a High Sensitivity DNA kit (Agilent, 5067-4626). Libraries were pooled and sequenced on a NovaSeq 6000 instrument at the Flow Cytometry & Single Cell Core Facility at the University of Copenhagen, Denmark.

### Data Analysis

Flow cytometry data was analyzed using Flow Jo v9.8.3 or v10. In lymphoid organs, γδT17 cells were gated as depicted in Figure S1A and in the brain they were identified as CD11b^−^CD45^+^CD3^+^CD4^−^TCRβ^−^TCRγδ^+^CD44^+^CD27^−^. All graphs not depicting single-cell data were generated and analyzed using Prism v8 or v10. Details for statistical tests are shown in each individual figure legend. For the scRNA-seq data, sequencing quality control and demultiplexing was carried out by the Flow Cytometry & Single Cell Core Facility at the University of Copenhagen. Reads were filtered and aligned to the GRCm39-2024-A reference genome using CellRanger v8.0.0. Seurat v5.1.0 was used for additional quality control and data analysis. Datasets were preprocessed individually; HTODemux was used to demultiplex hashed biological replicates, remove doublets and non-labeled cells. Cells with more than 5% mitochondrial counts and aberrantly low or high number of RNA molecules and genes were also excluded. In addition, ribosomal genes were removed. Afterwards, datasets were merged (e18 with newborn thymus; d0 control with d11 EAE spleen). The data was Log Normalized, and the effect of cell cycle variance was regressed out. Thymic and splenic datasets were integrated separately using Canonical Correlation Analysis (CCA). In the EAE data, we found contaminating myeloid cells (identified based on *Spi1*) and other non-T cells based on lack of *Cd3e* expression, and these were removed from the pool of cells. Then, Vγ4 and Vγ6 cells were subsetted based on the condition that cells express at least one transcript of *Trgv4* or *Trgv6*, respectively, and do not express any transcripts from the other chain. After subsetting, the data was re-scaled and re-integrated. Afterwards, unsupervised clustering was run with default parameters for the thymus, spleen Vγ4 and Vγ6 datasets. Cluster identities were found running Seurat’s “FindAllMarkers” function. Differential expression analysis was done using “FindMarkers”. In all cases, only genes expressed in at least 25% of cells and with a log_2_FC of ≥ 1 and an adjusted p-value of <0.05 (determined using a Wilcoxon Rank Sum test) were considered significant.

## Supplementary Figures

**Figure S1.**
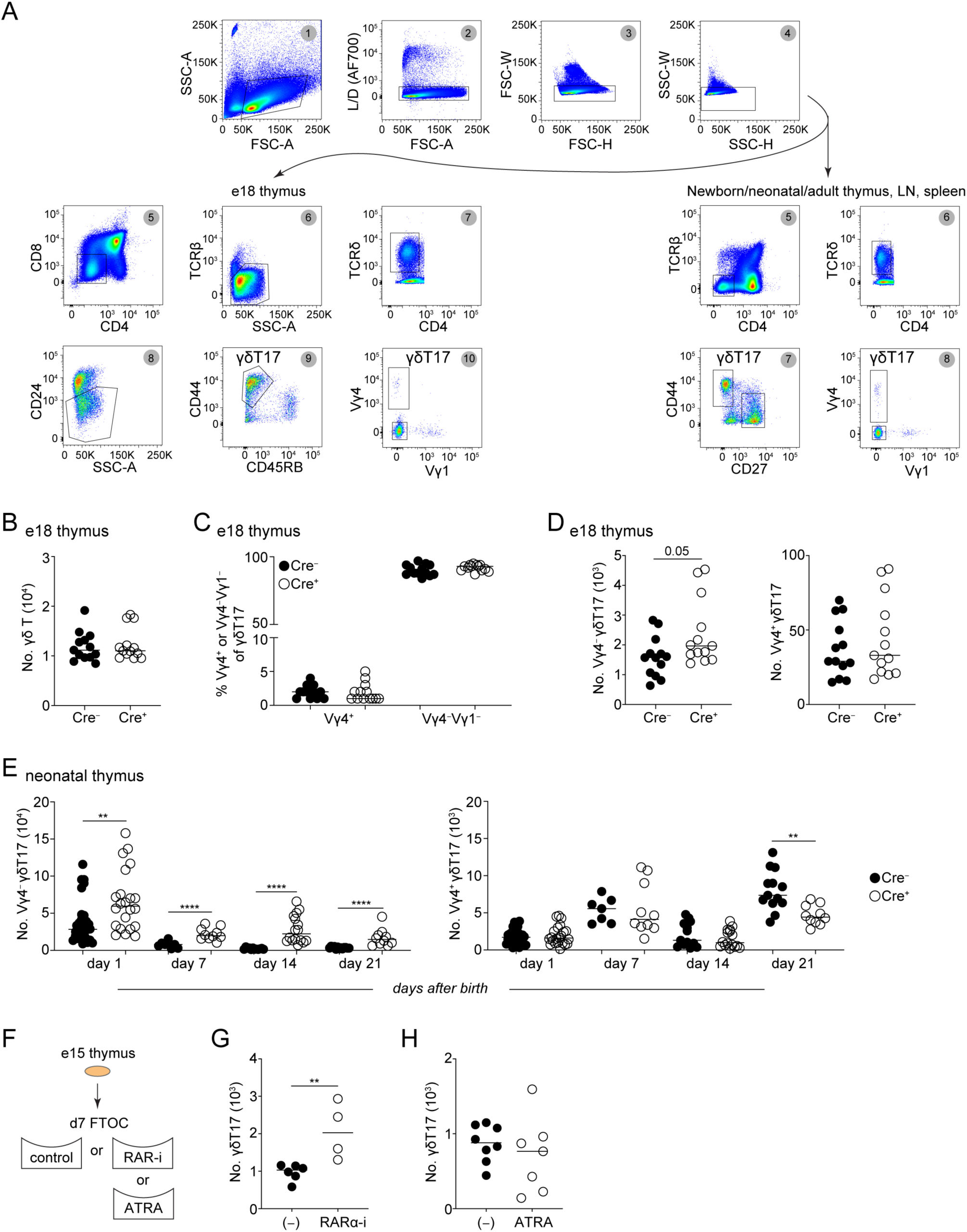
**RA limits the development of γδT17 cells.** (**A**) Sequential gating for the identification and enumeration of γδT17 cell populations in the embryonic thymus (left) and in all other neonatal and adult organs (right). (**B**) Numbers of total Cre^−^ and Cre^+^ γδ T cells at e18. (**C**) Frequency of Vγ4^+^ and Vγ4^−^Vγ1^−^ cells within the γδT17 population at e18. (**D**) Numbers of Vγ4^−^ (left) and Vγ4^+^ (right) cells within the γδT17 population at e18. (**E**) Numbers of Vγ4^−^ (left) and Vγ4^+^ (right) cells within the γδT17 population in the thymus at the indicated days after birth. (**F**) Schematic representation of the fetal thymic organ cultures (FTOCs). (**G-H**) Numbers of γδT17 cells in FTOCs that were treated with 8 μM RARα inhibitor (RARα-i) (**G**) or 160 nM of ATRA (**H**). In all graphs, each symbol represents one mouse (**B-E**) or 2-3 thymic lobes pooled (**G-H**); data are pool of three (**B-E**) or two (**G-H**) experiments; p-values were determined using Mann–Whitney test. **P < 0.01; ***P < 0.001.

**Figure S2.**
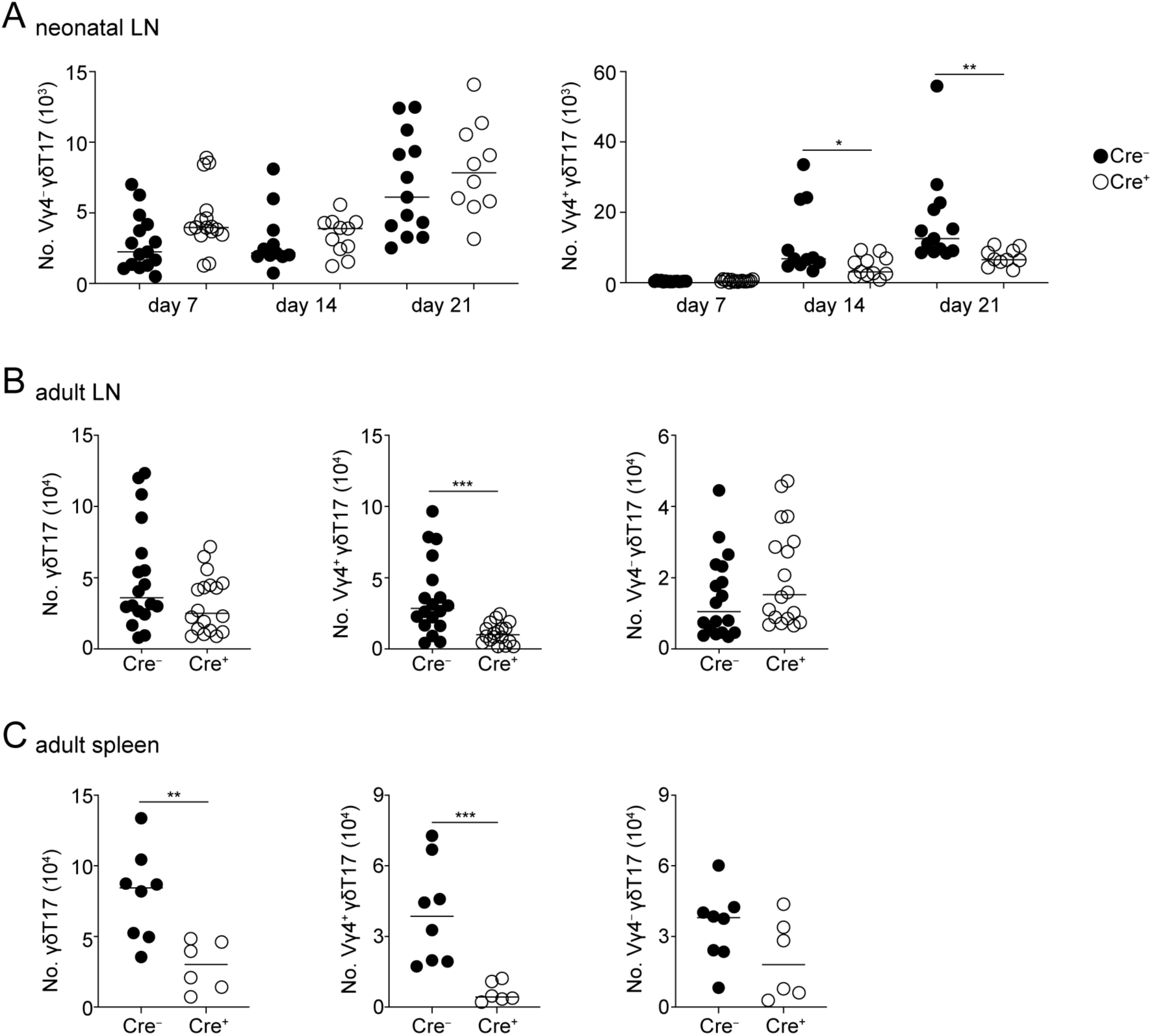
**RA regulates the numbers of γδT17 cells in neonatal and adult life.** (**A**) Numbers of Vγ4^−^ (left) and Vγ4^+^ (right) γδT17 cells in the LNs at the indicated days after birth. (**B-C**) Numbers of total γδT17 (left), Vγ4^+^ γδT17 (middle) and Vγ4^−^ γδT17 (right) cells in the LN and spleen (**C**) of adult Cre^−^ and Cre+ mice. In all graphs, each symbol represents one mouse; data are pool of three experiments; p-values were determined using Mann–Whitney test. *P < 0.05; **P < 0.01; ***P < 0.001.

**Figure S3.**
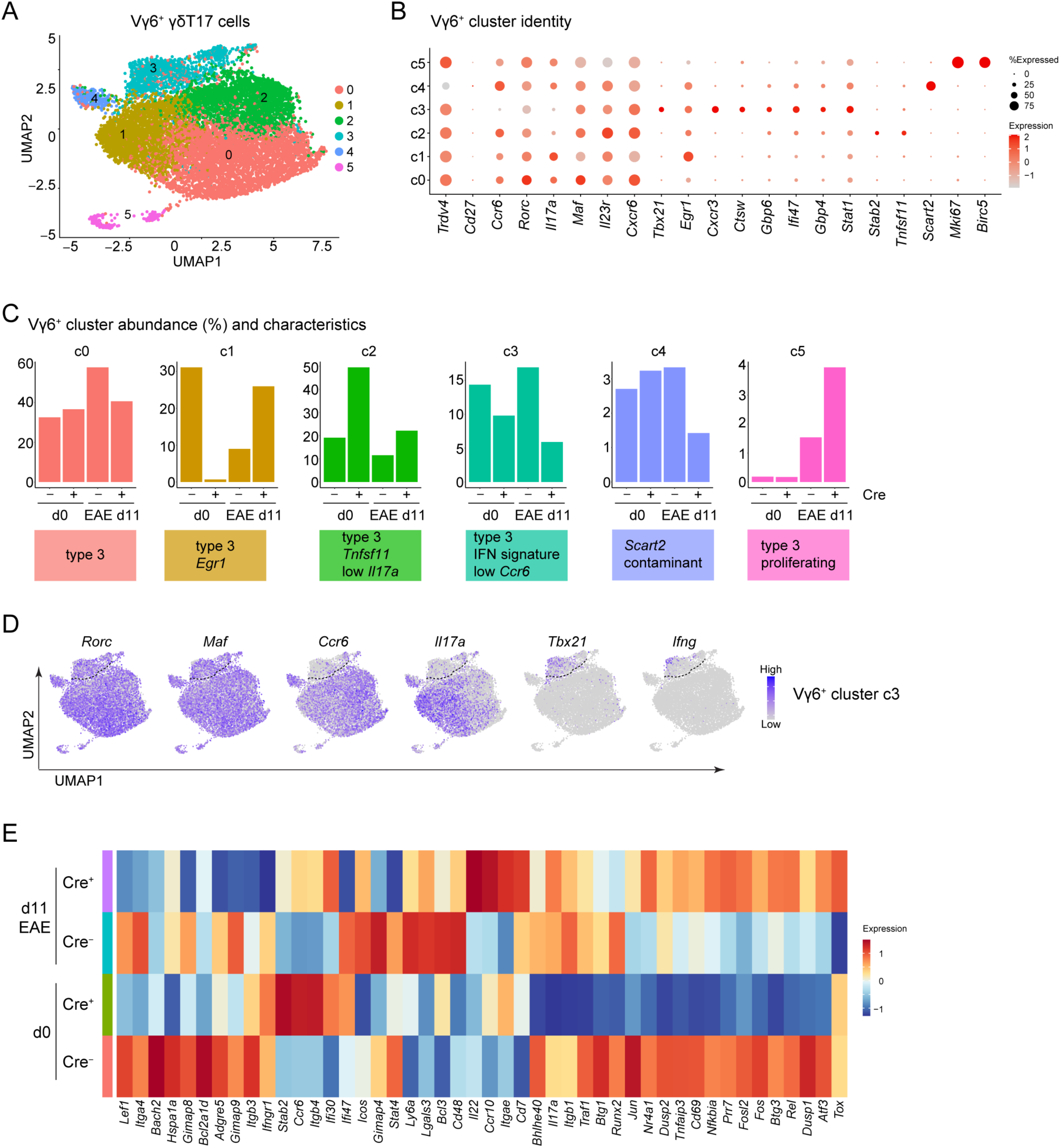
**RA induces activation, survival and interferon gene signatures in inflammation.** (**A**) UMAP of Vγ6^+^ γδT17 clusters, integrated from the spleen of Cre^−^ and Cre^+^ mice before (d0) and 11 days (d11) after MOG_35-55_ immunization. (**B**) Dot plot showing the scaled expression of selected genes across clusters, with red representing higher expression. Dot size indicates the percentage of cells within each cluster expressing the corresponding gene. (**C**) Cluster characterization and abundance within each genotype and timepoint, shown as a fraction of all Vγ6^+^ cells. (**D**) Feature plots illustrating the distribution and expression levels of *Rorc*, *Maf*, *Ccr6*, *Il17a*, *Tbx21* and *Ifng*. A line is drawn to delineate the IFN signature cluster c3 from the others. (**E**) Heatmap representation of selected DEGs between Cre^−^ and Cre^+^ cells for each EAE timepoint. Colors indicate scaled gene expression levels, with red representing higher expression.

**Figure S4.**
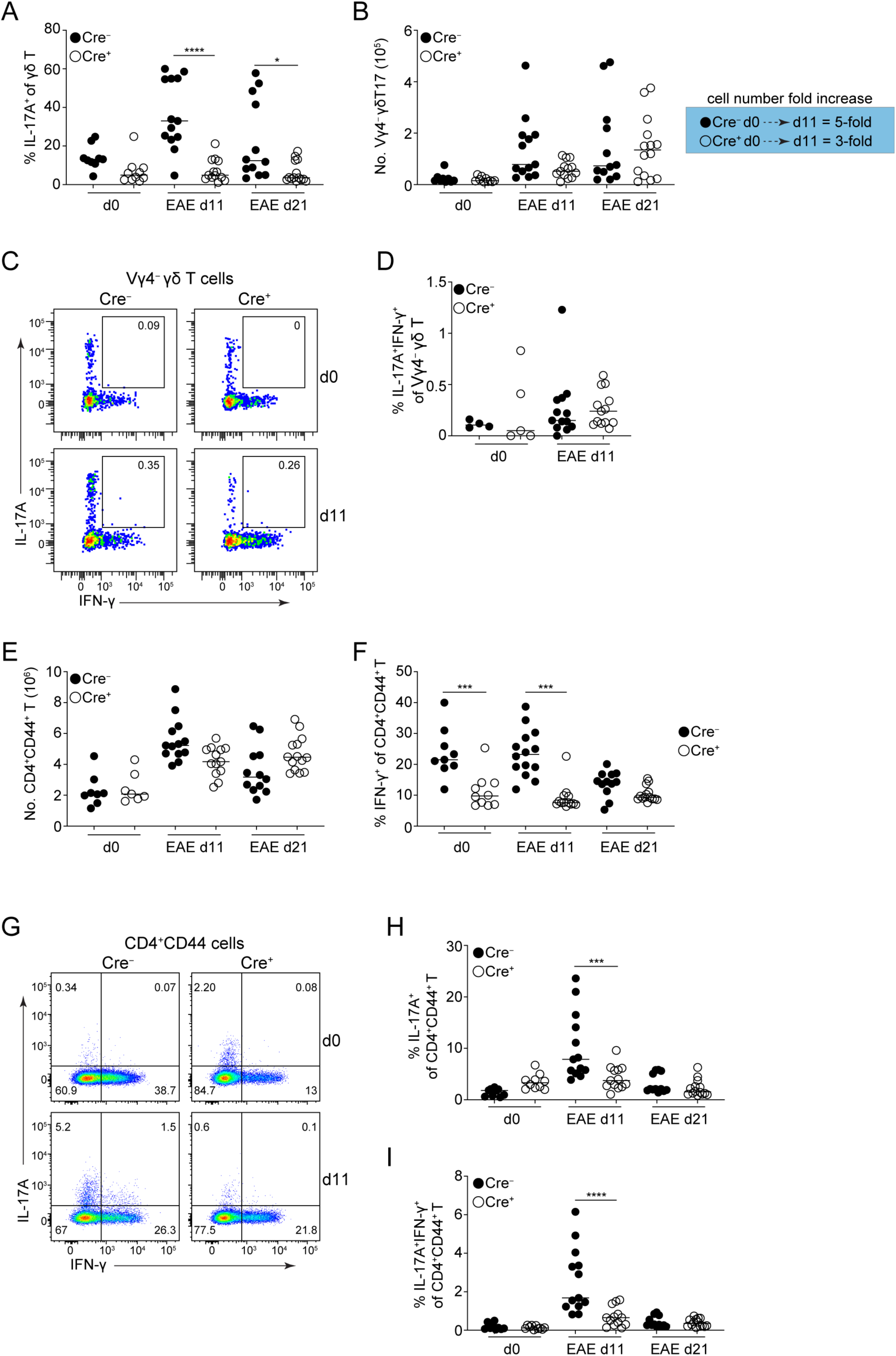
**γδ and CD4 T cell responses during EAE.** (**A**) Frequency of IL-17A^+^ γδ T cells in the spleen at d0, d11 and d21 of EAE. (**B**) Number of Vγ4^−^ cells within the γδT17 population in the spleen at different timepoints of EAE. On the right, the fold increase in cell numbers from d0 to d11 is displayed for each genotype. (**C**) Representative FACS plots for IL-17A and IFN-γ co-production in the spleen by Vγ4^−^ γδ T cells. (**D**) Frequency of IL- 17A^+^IFN-γ^+^ cells within the Vγ4^−^ γδ T cell population. (**E**) Number of splenic CD4^+^CD44^+^ T cells at d0, d11 and d21 of EAE. (**F**) Frequency of IFN-γ^+^ cells within the CD4^+^CD44^+^ T cell population at different timepoints of EAE. (**G**) Representative FACS plots for IL-17A and IFN-γ production in the spleen by activated (CD44^+^) Cre^−^ and Cre^+^ CD4^+^ T cells at d0 and d11. (**H-I**) Frequency of IL-17A^+^ (**H**) and IL-17A^+^IFN-γ^+^ (**I**) cells within the CD4^+^CD44^+^ T cell population at different EAE timepoints. For all plots, each symbol represents a mouse and the line the median; data are pool of three experiments; p-values determined using One Way ANOVA with Tukey’s multiple comparisons test. *P < 0.05; ***P < 0.001; ****P < 0.0001.

**Figure S5.**
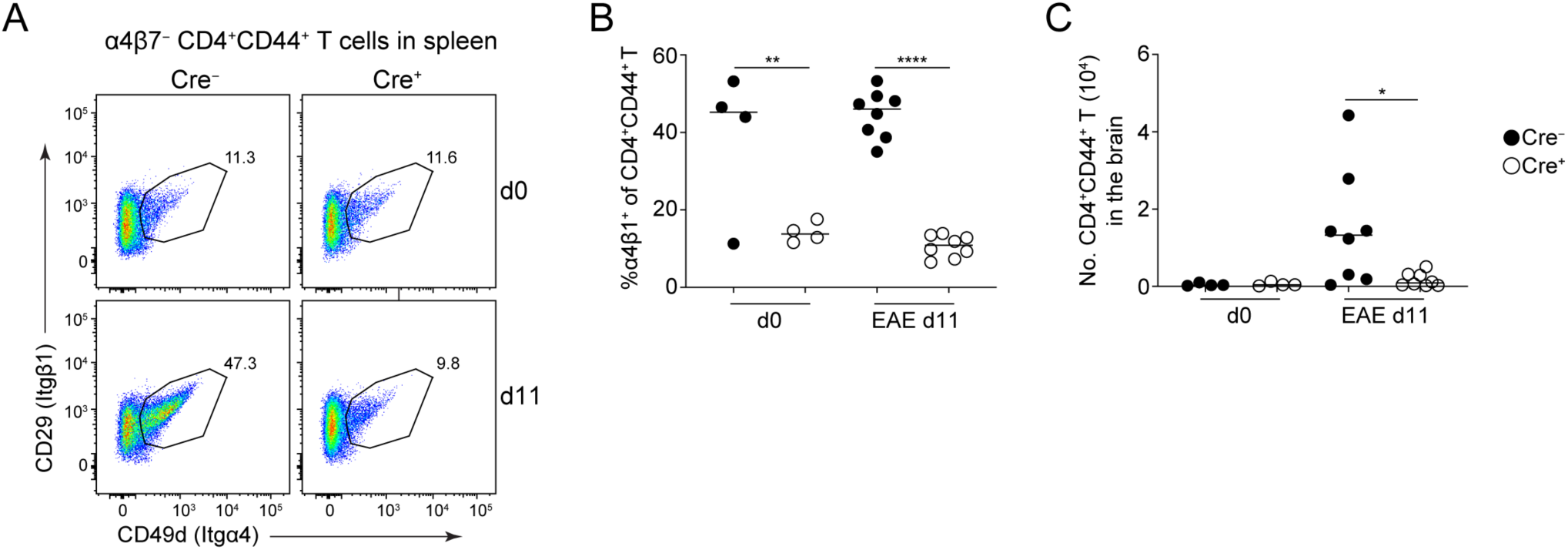
**RA is required for α4β1 expression and cell infiltration to the brain.** (**A**) Representative FACS plots of α4β1 integrin expression on splenic CD4^+^CD44^+^ T cells at d0 and d11 of EAE (cells were pre-gated as α4β7^−^). (**B**) Frequency of α4β1^+^ cells within the CD4^+^CD44^+^ T cell population. (**C**) Numbers of CD4^+^CD44^+^ T cells in the brain of Cre^−^ and Cre^+^ at d0 and d11 of EAE. In all graphs, each symbol represents one mouse; data are pool of two experiments; p-values determined using One Way ANOVA with Tukey’s multiple comparisons test. *P < 0.05; **P < 0.01; ****P < 0.0001.

## References

1. E. S. Edholm, M. Banach, J. Robert, Evolution of innate-like T cells and their selection by MHC class I-like molecules. Immunogenetics 68, 525–536 (2016).

2. M. Hirano et al., Evolutionary implications of a third lymphocyte lineage in lampreys. Nature 501, 435–438 (2013).

3. R. Agerholm, V. Bekiaris, Evolved to protect, designed to destroy: IL-17-producing gammadelta T cells in infection, inflammation, and cancer. Eur J Immunol, (2021).

4. J. C. Ribot et al., CD27 is a thymic determinant of the balance between interferon- gamma- and interleukin 17-producing gammadelta T cell subsets. Nat Immunol 10, 427–436 (2009).

5. G. Turchinovich, A. C. Hayday, Skint-1 identifies a common molecular mechanism for the development of interferon-gamma-secreting versus interleukin-17-secreting gammadelta T cells. Immunity 35, 59–68 (2011).

6. N. Sumaria, C. L. Grandjean, B. Silva-Santos, D. J. Pennington, Strong TCRgammadelta Signaling Prohibits Thymic Development of IL-17A-Secreting gammadelta T Cells. Cell reports 19, 2469–2476 (2017).

7. M. Munoz-Ruiz et al., TCR signal strength controls thymic differentiation of discrete proinflammatory gammadelta T cell subsets. Nat Immunol 17, 721–727 (2016).

8. M. K. Zuberbuehler et al., The transcription factor c-Maf is essential for the commitment of IL-17-producing gammadelta T cells. Nat Immunol 20, 73–85 (2019).

9. I. Prinz, B. Silva-Santos, D. J. Pennington, Functional development of gammadelta T cells. Eur J Immunol 43, 1988–1994 (2013).

10. D. Kadekar et al., The neonatal microenvironment programs innate gammadelta T cells through the transcription factor STAT5. J Clin Invest 130, 2496–2508 (2020).

11. J. Rizk et al., The cIAP ubiquitin ligases sustain type 3 gammadelta T cells and ILC during aging to promote barrier immunity. J Exp Med 220, (2023).

12. B. S. Sheridan et al., gammadelta T cells exhibit multifunctional and protective memory in intestinal tissues. Immunity 39, 184–195 (2013).

13. A. C. Kohlgruber et al., gammadelta T cells producing interleukin-17A regulate adipose regulatory T cell homeostasis and thermogenesis. Nat Immunol 19, 464–474 (2018).

14. A. O. Mann et al., IL-17A-producing gammadeltaT cells promote muscle regeneration in a microbiota-dependent manner. J Exp Med 219, (2022).

15. 15. M. Ribeiro, et al., Meningeal gammadelta T cell-derived IL-17 controls synaptic plasticity and short-term memory. *Sci Immunol* 4, (2019).

16. K. Alves de Lima et al., Meningeal gammadelta T cells regulate anxiety-like behavior via IL-17a signaling in neurons. Nat Immunol 21, 1421–1429 (2020).

17. M. N. Erkelens, R. E. Mebius, Retinoic Acid and Immune Homeostasis: A Balancing Act. Trends Immunol 38, 168–180 (2017).

18. D. N. D’Ambrosio, R. D. Clugston, W. S. Blaner, Vitamin A metabolism: an update. Nutrients 3, 63–103 (2011).

19. J. S. Steinhoff, A. Lass, M. Schupp, Biological Functions of RBP4 and Its Relevance for Human Diseases. Front Physiol 12, 659977 (2021).

20. N. B. Ghyselinck, G. Duester, Retinoic acid signaling pathways. Development 146, (2019).

21. J. A. Hall, J. R. Grainger, S. P. Spencer, Y. Belkaid, The role of retinoic acid in tolerance and immunity. Immunity 35, 13–22 (2011).

22. G. Duester, Retinoic acid synthesis and signaling during early organogenesis. Cell 134, 921–931 (2008).

23. C. O’Connor, P. Varshosaz, A. R. Moise, Mechanisms of Feedback Regulation of Vitamin A Metabolism. Nutrients 14, (2022).

24. M. R. Conserva, L. Anelli, A. Zagaria, G. Specchia, F. Albano, The Pleiotropic Role of Retinoic Acid/Retinoic Acid Receptors Signaling: From Vitamin A Metabolism to Gene Rearrangements in Acute Promyelocytic Leukemia. Int J Mol Sci 20, (2019).

25. N. R. de Almeida, M. Conda-Sheridan, A review of the molecular design and biological activities of RXR agonists. Med Res Rev 39, 1372–1397 (2019).

26. S. A. van de Pavert et al., Chemokine CXCL13 is essential for lymph node initiation and is induced by retinoic acid and neuronal stimulation. Nat Immunol 10, 1193–1199 (2009).

27. S. A. van de Pavert et al., Maternal retinoids control type 3 innate lymphoid cells and set the offspring immunity. Nature 508, 123–127 (2014).

28. W. W. Agace, E. K. Persson, How vitamin A metabolizing dendritic cells are generated in the gut mucosa. Trends Immunol 33, 42–48 (2012).

29. K. M. Luda et al., IRF8 Transcription-Factor-Dependent Classical Dendritic Cells Are Essential for Intestinal T Cell Homeostasis. Immunity 44, 860–874 (2016).

30. M. H. Kim, E. J. Taparowsky, C. H. Kim, Retinoic Acid Differentially Regulates the Migration of Innate Lymphoid Cell Subsets to the Gut. Immunity 43, 107–119 (2015).

31. J. L. Coombes et al., A functionally specialized population of mucosal CD103+ DCs induces Foxp3+ regulatory T cells via a TGF-beta and retinoic acid-dependent mechanism. J Exp Med 204, 1757–1764 (2007).

32. C. M. Sun et al., Small intestine lamina propria dendritic cells promote de novo generation of Foxp3 T reg cells via retinoic acid. J Exp Med 204, 1775–1785 (2007).

33. D. Mucida et al., Reciprocal TH17 and regulatory T cell differentiation mediated by retinoic acid. Science 317, 256–260 (2007).

34. H. R. Cha et al., Downregulation of Th17 cells in the small intestine by disruption of gut flora in the absence of retinoic acid. J Immunol 184, 6799–6806 (2010).

35. J. A. Hall et al., Essential role for retinoic acid in the promotion of CD4(+) T cell effector responses via retinoic acid receptor alpha. Immunity 34, 435–447 (2011).

36. C. C. Brown et al., Retinoic acid is essential for Th1 cell lineage stability and prevents transition to a Th17 cell program. Immunity 42, 499–511 (2015).

37. B. S. Reis, D. P. Hoytema van Konijnenburg, S. I. Grivennikov, D. Mucida, Transcription factor T-bet regulates intraepithelial lymphocyte functional maturation. Immunity 41, 244–256 (2014).

38. B. S. Reis, A. Rogoz, F. A. Costa-Pinto, I. Taniuchi, D. Mucida, Mutual expression of the transcription factors Runx3 and ThPOK regulates intestinal CD4(+) T cell immunity. Nat Immunol 14, 271–280 (2013).

39. A. Obers et al., Retinoic acid and TGF-beta orchestrate organ-specific programs of tissue residency. Immunity 57, 2615–2633 e2610 (2024).

40. Z. Qiu et al., Retinoic acid signaling during priming licenses intestinal CD103+ CD8 TRM cell differentiation. J Exp Med 220, (2023).

41. M. Raverdeau, C. J. Breen, A. Misiak, K. H. Mills, Retinoic acid suppresses IL-17 production and pathogenic activity of gammadelta T cells in CNS autoimmunity. Immunol Cell Biol 94, 763–773 (2016).

42. L. A. Mielke et al., Retinoic acid expression associates with enhanced IL-22 production by gammadelta T cells and innate lymphoid cells and attenuation of intestinal inflammation. J Exp Med 210, 1117–1124 (2013).

43. M. Lochner et al., In vivo equilibrium of proinflammatory IL-17+ and regulatory IL-10+ Foxp3+ RORgamma t+ T cells. J Exp Med 205, 1381–1393 (2008).

44. F. Rajaii, Z. T. Bitzer, Q. Xu, S. Sockanathan, Expression of the dominant negative retinoid receptor, RAR403, alters telencephalic progenitor proliferation, survival, and cell fate specification. Dev Biol 316, 371–382 (2008).

45. J. S. Heilig, S. Tonegawa, Diversity of murine gamma genes and expression in fetal and adult T lymphocytes. Nature 322, 836–840 (1986).

46. C. M. Carlson et al., Kruppel-like factor 2 regulates thymocyte and T-cell migration. Nature 442, 299–302 (2006).

47. S. Filen, R. Lahesmaa, GIMAP Proteins in T-Lymphocytes. J Signal Transduct 2010, 268589 (2010).

48. N. Sumaria, S. Martin, D. J. Pennington, Constrained TCRgammadelta-associated Syk activity engages PI3K to facilitate thymic development of IL-17A-secreting gammadelta T cells. Sci Signal 14, (2021).

49. F. Coffey et al., The TCR ligand-inducible expression of CD73 marks gammadelta lineage commitment and a metastable intermediate in effector specification. J Exp Med 211, 329–343 (2014).

50. A. M. McGinley et al., Interleukin-17A Serves a Priming Role in Autoimmunity by Recruiting IL-1beta-Producing Myeloid Cells that Promote Pathogenic T Cells. Immunity 52, 342–356 e346 (2020).

51. C. E. Sutton et al., Interleukin-1 and IL-23 induce innate IL-17 production from gammadelta T cells, amplifying Th17 responses and autoimmunity. Immunity 31, 331–341 (2009).

52. Ivanov, II et al., The orphan nuclear receptor RORgammat directs the differentiation program of proinflammatory IL-17+ T helper cells. Cell 126, 1121–1133 (2006).

53. K. H. Johansen et al., A CRISPR screen targeting PI3K effectors identifies RASA3 as a negative regulator of LFA-1-mediated adhesion in T cells. Sci Signal 15, eabl9169 (2022).

54. B. Wu et al., RAS P21 Protein Activator 3 (RASA3) Specifically Promotes Pathogenic T Helper 17 Cell Generation by Repressing T-Helper-2-Cell-Biased Programs. Immunity 49, 886–898 e885 (2018).

55. E. P. Bowman et al., Regulation of chemotactic and proadhesive responses to chemoattractant receptors by RGS (regulator of G-protein signaling) family members. J Biol Chem 273, 28040–28048 (1998).

56. K. G. Haanstra et al., Antagonizing the alpha4beta1 integrin, but not alpha4beta7, inhibits leukocytic infiltration of the central nervous system in rhesus monkey experimental autoimmune encephalomyelitis. J Immunol 190, 1961–1973 (2013).

57. B. E. Theien et al., Discordant effects of anti-VLA-4 treatment before and after onset of relapsing experimental autoimmune encephalomyelitis. J Clin Invest 107, 995–1006 (2001).

58. G. P. Rice, H. P. Hartung, P. A. Calabresi, Anti-alpha4 integrin therapy for multiple sclerosis: mechanisms and rationale. Neurology 64, 1336–1342 (2005).

59. T. A. Yednock et al., Prevention of experimental autoimmune encephalomyelitis by antibodies against alpha 4 beta 1 integrin. Nature 356, 63–66 (1992).

60. J. L. Baron, J. A. Madri, N. H. Ruddle, G. Hashim, C. A. Janeway, Jr., Surface expression of alpha 4 integrin by CD4 T cells is required for their entry into brain parenchyma. J Exp Med 177, 57–68 (1993).

61. N. A. Spidale et al., Interleukin-17-Producing gammadelta T Cells Originate from SOX13(+) Progenitors that Are Independent of gammadeltaTCR Signaling. Immunity 49, 857–872 e855 (2018).

62. Y. Ohoka, A. Yokota, H. Takeuchi, N. Maeda, M. Iwata, Retinoic acid-induced CCR9 expression requires transient TCR stimulation and cooperativity between NFATc2 and the retinoic acid receptor/retinoid X receptor complex. J Immunol 186, 733–744 (2011).

63. A. Larange et al., A regulatory circuit controlled by extranuclear and nuclear retinoic acid receptor alpha determines T cell activation and function. Immunity 56, 2054–2069 e2010 (2023).

64. N. Noy, Between death and survival: retinoic acid in regulation of apoptosis. Annu Rev Nutr 30, 201–217 (2010).

65. F. Da Silva et al., Retinoic acid signaling is directly activated in cardiomyocytes and protects mouse hearts from apoptosis after myocardial infarction. eLife 10, (2021).

66. M. Iwata, M. Mukai, Y. Nakai, R. Iseki, Retinoic acids inhibit activation-induced apoptosis in T cell hybridomas and thymocytes. J Immunol 149, 3302–3308 (1992).

67. Y. Yang, M. S. Vacchio, J. D. Ashwell, 9-cis-retinoic acid inhibits activation-driven T-cell apoptosis: implications for retinoid X receptor involvement in thymocyte development. Proc Natl Acad Sci U S A 90, 6170–6174 (1993).

68. F. Petermann et al., gammadelta T cells enhance autoimmunity by restraining regulatory T cell responses via an interleukin-23-dependent mechanism. Immunity 33, 351–363 (2010).

69. C. A. Dendrou, L. Fugger, M. A. Friese, Immunopathology of multiple sclerosis. Nat Rev Immunol 15, 545–558 (2015).

70. M. Hiltensperger, T. Korn, The Interleukin (IL)-23/T helper (Th)17 Axis in Experimental Autoimmune Encephalomyelitis and Multiple Sclerosis. Cold Spring Harb Perspect Med 8, (2018).

71. L. Schirmer, V. Rothhammer, B. Hemmer, T. Korn, Enriched CD161high CCR6+ gammadelta T cells in the cerebrospinal fluid of patients with multiple sclerosis. JAMA Neurol 70, 345–351 (2013).

72. J. van Langelaar et al., T helper 17.1 cells associate with multiple sclerosis disease activity: perspectives for early intervention. Brain 141, 1334–1349 (2018).

73. E. Havrdova et al., Activity of secukinumab, an anti-IL-17A antibody, on brain lesions in RRMS: results from a randomized, proof-of-concept study. J Neurol 263, 1287–1295 (2016).

74. C. Luckel et al., IL-17(+) CD8(+) T cell suppression by dimethyl fumarate associates with clinical response in multiple sclerosis. Nature communications 10, 5722 (2019).

75. V. Cipollini, J. Anrather, F. Orzi, C. Iadecola, Th17 and Cognitive Impairment: Possible Mechanisms of Action. Front Neuroanat 13, 95 (2019).

76. V. Brucklacher-Waldert et al., Tbet or Continued RORgammat Expression Is Not Required for Th17-Associated Immunopathology. J Immunol 196, 4893–4904 (2016).

77. K. Hirota et al., Fate mapping of IL-17-producing T cells in inflammatory responses. Nat Immunol 12, 255–263 (2011).

78. C. Klemann et al., Synthetic retinoid AM80 inhibits Th17 cells and ameliorates experimental autoimmune encephalomyelitis. Am J Pathol 174, 2234–2245 (2009).

